# Reshaping epigenomic landscapes in facilitating the speciation of bread wheat

**DOI:** 10.1101/2024.12.26.630455

**Authors:** Zhaoheng Zhang, Xuelei Lin, Jingjing Yue, Yongxin Xu, Lingfeng Miao, Wenqiang Tang, Weilong Guo, Jun Xiao

## Abstract

Polyploidization is a driving force of wheat evolution and speciation, yet its impact on epigenetic regulation and gene expression remains unclear.

Here, we constructed a high-resolution epigenetic landscape across leaves, spikes, and roots of hexaploidy wheat and its tetraploid and diploid relatives. Inter-species stable-expression genes exhibited conserved amino acid sequences under strong purifying selection, while dynamic-expression genes were linked to species-specific adaptation. During hexaploidization, dominant D-subgenome homoeolog expression was suppressed via reduced activating epigenetic signals, converging expression with the A and B subgenomes. Proximal chromatin regions near genes were more stable, whereas distal regions, particularly enhancer-like elements mediated by H3K27ac and H3K4me3, exhibit higher dynamism. Sequence variations in these enhancers lead to differential gene regulation, influencing traits such as spike development. For instance, the two haplotypes of dCRE region of *TaDEP-B1* resulted in significant differences in its expression and spikelet numbers. We also observed a coevolution of transcription factors and their binding sites, particularly within the expanded ERF family, which regulates spike morphology.

This study highlights the interplay between sequence variation and epigenetic modifications in shaping transcriptional regulation during wheat speciation, offering valuable insights for genetic improvement.

## Introduction

Polyploidy, characterized by the presence of two or more sets of chromosomes in an organism or cell, is a frequent occurrence in angiosperms such as wheat, cotton, and rapeseed (Yang *et al*., 2016; Wang, M *et al*., 2017; Ramírez-González *et al*., 2018). Polyploidization plays a vital role in plant evolution by creating new phenotypes, enhancing species diversity, and improving adaptability to environmental factors (Mable *et al*., 2011; Van de Peer *et al*., 2017). It also has potential applications in agriculture, such as increasing drought resistance, insect resistance, and biomass (Osborn *et al*., 2003). However, the genetic redundancy resulting from polyploidization can induce changes in gene expression through both genetic and epigenetic regulation, leading to the re-organization of the genome and regulatory networks (Song & Chen, 2015; Van de Peer *et al*., 2017). Although substantial advancements have been made in studying the epigenome in diploid plants, particularly in *Arabidopsis* and rice. The integrated mechanisms governing the epigenome and transcriptome interactions in this context remain undisclosed in polyploid, presenting a complex challenge in coordinating subgenomes within individual cell nuclei (Yu *et al*., 2021).

Bread wheat, one of the important domesticated crops, has been a staple food for thousands of years, providing a significant portion of daily consumed calories and proteins (Shiferaw *et al*., 2013). It originated from polyploidization events within the *Triticum* and *Aegilops* genera, resulting in the allohexaploid species with A, B, and D subgenomes (Consortium *et al*., 2014; Xiao *et al*., 2022). The first allotetraploidization, which occurred approximately 0.5-3 million years ago, involved hybridization between *Triticum urartu* (AA) and an undiscovered or extinct species closely related to *Aegilops speltoides* (SS). The second allohexaploidization, between domesticated tetraploid wheat (AABB) and *Aegilops tauschii* (DD) approximately 8,000-10,000 years ago, resulted in the formation of bread wheat (Consortium *et al*., 2014; Zhang *et al*., 2014). The fusion of genomes from different environments during polyploidization expanded the adaptability of bread wheat (Dubcovsky & Dvorak, 2007; Akhunov *et al*., 2010; Van de Peer *et al*., 2017). Human selection for agronomic traits further contributed to its global adaptations, while maintaining some variation in specific characteristics to optimize fitness for different growth regions (Zhou *et al*., 2020). Concurrently, diploid and tetraploid ancestral species serve as valuable genetic resources for enhancing wheat yield (Katamadze *et al*., 2023). Currently, scientists have harnessed synthetic wheats (SHWs), which are artificial hybrids created by crossing *Triticum turgidum* (BBAA) with *Aegilops tauschii* (DD), to produce varieties characterized by larger spikes and grains, thereby significantly contributing to the improvement of wheat spike development (Yang *et al*., 2009). The hybridization that results in synthetic wheats has yielded wheat varieties with increased tolerance to biotic and abiotic stresses and higher yield (Ogbonnaya *et al*., 2007; Lopes & Reynolds, 2011; Ogbonnaya *et al*., 2013). Therefore, elucidating the regulatory mechanisms underlying the traits of diploid and tetraploid wheat will facilitate the identification of novel genes and superior alleles for wheat improvement.

Gene expression in hexaploid wheat involves the coordination of transcription and epigenetic regulation among different subgenomes. Highly expressed genes exhibit preferential active histone modification marks, such as H3K4me3, H3K9ac, H3K27ac, and H3K36me3, while tissue-specific expressed genes are likely associated with H3K27me3 (Wang, M *et al*., 2021). In the face of “genomic shock” caused by polyploidization, significant changes occur in chromatin dynamics (Liu *et al*., 2021; Yuan *et al*., 2022) and DNA methylation in wheat (Yuan *et al*., 2020; Miao *et al*., 2024). Intensity of H3K27me2 in wheat increases with the level of ploidy, which contributes to silence euchromatic transposons to maintain genome stability and modify genetic recombination landscapes (Liu et al., 2021). Genome merging and separation in polyploid wheat also result in dynamic and reversible changes in DNA methylation to regulate gene expression and transposon activity (Yuan et al., 2020). The open chromatin enrichment and network Hi-C (OCEAN-C) in different ploidy wheat revealed the regulation of gene expression by chromatin interaction changes caused by genomic structural variation (Yuan et al., 2022). In *Brassica napus*, activating epigenetic marks have been observed to drive subgenome dominance (Zhang *et al*., 2021). Although this phenomenon has not been observed in wheat, the expression bias of homoeologs and the divergence in subgenome functions in hexaploid wheat are associated with epigenetic modifications (Wang, M *et al*., 2021). Moreover, recent studies have unveiled a complex regulatory mechanism mediated by a series of epigenetic marks during the development of grain and spikes in bread wheat (Zhao *et al*., 2023; Lin *et al*., 2024; Zhao *et al*., 2024). However, the holistic dynamics regarding the influence of chromatin state defined by multiple histone modifications on wheat subgenomes at different stages during hexaploid wheat species formation remain unexplored. This is crucial for comprehending both the shared aspects of polyploid formation and the distinctive characteristics of hexaploid wheat.

In this study, we generated high-resolution profiles of core histone modifications in hexaploid wheat and its tetraploid and diploid relatives, uncovering the critical role of epigenetic modifications in driving transcriptional dynamics during hexaploidy wheat speciation. Our findings highlight the dynamic contributions of distal enhancer elements to phenotypic variation and the co-evolution of transcription factors (TFs) with their binding sites throughout wheat speciation. This study underscores the synergistic interplay between DNA sequence variations and chromatin modification changes, offering deeper insights into the regulatory mechanisms driving wheat evolution.

## Materials and Methods

### Plant materials and growth conditions

*T. urartu* (AA, *c.v.* G1812)*, Ae. tauschii* (DD, *c.v.* AL8/78)*, T. dicoccoides* (AABB, *c.v.* Zavitan)*, T. durum* (AABB, *c.v.* Svevo) and *T. aestivum* (AABBDD, *c.v.* KN9204) were used in this study. The germinated seeds were treated at 4 °C for 30 days. The seedlings were transplanted into soil and grown in the greenhouse at 22 °C/20 °C day/night, under long day conditions (16 h light/8 h dark). The leaves and roots (one leaf and one heart), spikes (lemma primordium stage, W3.25) were collected for the further experiments. *N. benthamiana* was grown in a greenhouse at 22□°C under a 16 h light and 8 h darkness photoperiod.

### RNA-seq, ATAC-seq and CUT&Tag experiment

Total RNA was extracted using HiPure Plant RNA Mini Kit (Magen, R4111-02). RNA-seq library construction and sequencing platform were the same as previous description (Zhao et al., 2023), by Annoroad Gene Technology.

ATAC-seq and CUT&Tag experiments followed the previously described (Zhao et al., 2023). Tn5 transposase was used and tagmentation assay was performaed following the manual (Vazyme, TD501-01). Libraries were purified with AMPure beads (Beckman, A63881) and sequenced using the Illumina Novaseq platform at Annoroad Gene Technology. Antibodies used for histone modifications are listed in **Table S9**.

### Data Preprocessing and Alignment

To ensure the high quality of the reads, an initial filtering process was performed using fastp v0.20.1 with parameter “--detect_adapter_for_pe”. This included adapter removal, trimming of low-quality bases, and reads filtering (Chen *et al*., 2018). Furthermore, FastQC v0.11.8 (http://www.bioinformatics.babraham.ac.uk/projects/fastqc/) was performed to ensure the high quality of reads.

For the alignment of the reads, two different tools were employed based on the sequencing methodology used. BWA-MEM v0.7.17 (Li & Durbin, 2010) with parameter “-M” (for ATAC-seq and CUT&Tag seq), while hisat2 v2.1.0 (Kim *et al*., 2019) with default parameters was used for RNA-seq data. The reference genome used *Triticum aestivum c.v.* Chinese Spring (IWGSC RefSeq v1.0, https://urgi.versailles.inra.fr/download/iwgsc/IWGSC_RefSeq_Assemblies/v1.0/).

The reads from AA (*T. urartu*), AABB (*T. dicoccoides* and *T. durum*), and DD (*Ae. tauschii*) were mapped to the A, AB, and D subgenomes of reference sequences of IWGSC RefSeq v1.0, respectively. The reference for gene models was IWGSC Annotation v1.1, and only high-confidence genes were considered in this study. The resulting alignments were converted to BAM format, sorted, and indexed using Samtools v1.4 (Danecek *et al*., 2021). For RNA-seq data aligned with hisat2, deduplication was not performed during the conversion from SAM to BAM format. while, for ATAC-seq and CUT&Tag data, the SAM files were further filtered using the samtools with “view -bS -F 1,804 -f 2 -q 30” command to remove low-quality mapped reads. Duplicates in the high-quality mapped reads were removed using Picard v2.23.3 (https://github.com/broadinstitute/picard). Two replicates’ BAM files were then merged using samtools. To normalize and visualize the individual and merged replicate datasets, the BAM files were converted to bigwig files using the bamCoverage tool provided by deepTools v3.3.0 (Ramírez *et al*., 2014). The normalization was performed using CPM (Counts per Million mapped reads).

### RNA-seq data analyses

RNA-seq libraries’ construction and sequencing platform were the same as previous description (Zhao et al., 2023), by Annoroad Gene Technology. To quantify the number of paired reads mapped to each gene, featureCounts v2.0.1 with the parameter “-p” was utilized (Ramírez et al., 2014). The resulting counts files were then used for differential expression analysis using DESeq2 v1.26.0 (Love *et al*., 2014), with a threshold of “|Log2 Fold Change| ≥ 0.75 and FDR≤0.05” to identify differentially expressed genes. The raw counts were normalized to TPM (Transcripts Per Kilobase Million) for gene expression quantification. For clustering and visualization purposes, mean TPM were obtained by merging three biological replicates. Gene TPM values were z-scaled and clustered using the *k*-means method, and visualization was done using the R package ComplexHeatmap v2.4.3 (Gu, 2022). Functional enrichment analysis was performed using the R package clusterProfiler v3.18.1 (Yu *et al*., 2012), with GO annotation files generated from IWGSC Annotation v1.1. If not specified the TPM were further normalized to log_2_(TPM+1) for visualization.

### CUT&Tag and ATAC-seq data analyses

Data processing and reads alignment were performed as previously described (Zhao et al., 2023). MACS2 v2.1.4 (Feng *et al*., 2012) was used for peak calling, with parameters “-p 1e-3” for H3K27ac, H3K4me3, and H2A.Z, and parameters “--broad – broad-cutoff 0.05” for H3K27me3 and H3K36me3. For ATAC-seq data, MACS2 was used with parameters “--cutoff-analysis --nomodel --shift -100 --extsize 200”. The resulting peaks were annotated using the ChIPseeker v1.26.2 R package and the “annotatePeak” function (Yu *et al*., 2015). Gene promoters were defined as 3.0 Kb upstream to downstream 1.0 Kb of the TSS

For quantification of ATAC-seq and CUT&Tag data, read counts under the reference peak and the normalized counts per million (DBA_SCORE_TMM_READS_EFFECTIVE_CPM) values were generated using the R package DiffBind v3.12 (Bressan *et al*., 2024). The CPM values of the reference peak were used to perform PCA and hierarchical clustering analysis. The raw peak counts were used as input to identify differentially accessible peaks and differentially marked peaks by histone modification using the R package DESeq2. The heat maps centred on peaks were created using computeMatrix and plotHeatmap from deepTools.

To calculate correlations between transcriptome and epigenome data, Z-scaled TPM values of genes and Z-scaled CPM values peaks annotated to promoter were calculated. Then PCCs were calculated in R v4.3.2.

### Chromatin states analysis

For CS analysis, chromHMM was used “BinarizeBam” and “LearnModel” commands were used for chromatin-state annotation (Ernst & Kellis, 2012). We used 5 CUT&Tag (H3K4me3, H3K27ac, H3K27me3, H3K36me3 and H2A.Z) data as input. Five to twenty-five chromatin-states were generated and we selected the 12-chromatin-states as the final chromatin states. The calculation method for chromatin conservation is to use the “segments-merge.bed” file outputted by “LearnModel” as input and compute the proportion of identical chromatin bases between samples using a window size of 2 Mb.

### Similarity analysis of chromatin states

The entire genome is segmented into 200 bp bins, and the similarity of the chromatin states in each bin between each pair of species (*T. urartu* vs *T. dicoccoides*, *T. dicoccoides* vs *T. durum*, *T. durum* vs *T. aestivum* and *Ae. tauschii* vs *T. aestivum*) was calculated based on the evolutionary history, referring to previous study (Zhang et al., 2021). Finally, the average similarity between all species was calculated for each bin.

### Analysis of Ka/Ks

To calculate Ka/Ks values of the homoeolog gene pairs, the protein sequences of each gene and the corresponding coding sequence (CDS) were obtained by annotating the gene. The information of homoeolog gene pairs across hexaploidy wheat and its tetraploid and diploid relatives download from TGT (http://wheat.cau.edu.cn/TGT/). The NG methods for estimating Ka and Ks and their references were used to calculate Ka/Ks values using the TBtools (Chen *et al*., 2020). Then we obtained the Ka/Ks values reported in the main text. Instances where the Ka/Ks value was infinite (due to a Ks value of 0, and a non-zero Ka) were changed to 10 as a comparatively large, non-infinite number, while negative and NaN values of Ka/Ks (due to missing information or zero Ka and Ks values, respectively) were considered as “NA” values and excluded from further analysis.

### Prediction of H3K27ac/H3K4me3-mediated distal regulatory sites

The distal regulatory regions annotation strategy was largely based on a previous study (Corces et al., 2018; Zhang et al., 2023). Genes within 0.5 Mb from a distal H3K27ac peak are considered candidate target genes. Then, we generated null model as correlations between randomly selected peaks and randomly selected genes on different chromosomes, and enabling us to compute mean and standard deviation of this null distribution. For each potential link, after calculate correlation between gene expression (TPM) and distal H3K27ac or H3K4me3 signal (CPM) in samples, we also compute *P*-values for the test correlations based on null model, then significantly pairs were selected as regulatory region-gene pairs.

### Luciferase reporter assay

For LUC analyses, full-length coding sequences of *WFZP* from KN9204 was cloned into PTF101 vector to generate the effector construct p35S: *WFZP*-GFP; About 2 Kb promoter fragment of *TaSPL14-A1* and *TaDUO-A1* were amplified and fused in-frame with the CP461-LUC vector to generate the reporter construct p*TaSPL14-A1*:LUC and p*TaDUO-A1*:LUC. The plasmids were transformed into *A. tumefaciens* GV3101.

To generate *TaDEP-B1*-mini35Spro:LUC construct, the genomic sequence of distal regulatory region was amplified and fused in-frame with the pMY155-mini35S vector to generate the reporter construct. Then, mini35Spro:LUC (as control) and *TaDEP-B1*-mini35Spro:LUC were transformed into *A. tumefaciens* GV3101.

Then these strains were injected into *N. benthamiana* leaves (6∼8 leaf stage) in different combinations with p19, which was used to suppress RNA silencing. Dual luciferase assay reagents (Promega; E1910) with the Renilla luciferase gene as an internal control were used for luciferase imaging. The Dual-Luciferase Reporter Assay System (Promega; E2940) was used to quantify fluorescence signals. Relative LUC activity was calculated by the ratio of LUC/REN. The relevant primers are listed in **Table S9**.

### Detection and enrichment analysis of transcription factor binding motifs

We detected the proximal (distance TSS -3000 to 1000) and distal (H3K27ac- and H3K4me3-mediated dCRE) ACR regions. For the subsequent motif analysis, we downloaded the position weight matrices of 619 plant motif from PlantTFdb (Jin *et al*., 2016). Similar to the alignment process, SNPs calling from *T. urartu, Ae. tauschii, T. dicoccoides* and *T. durum* genome assemblies were replaced into the Chinese spring genome as their respective reference genomes using the “FastaAlternateReferenceMaker” module of GATK v4.4.4 (McKenna *et al*., 2010) before the motif scanning was performed. Fimo v5.4.1 (Bailey *et al*., 2009) was used to scan for motifs within ACRs. The motifs identified in the proximal regulatory region of a gene are defined as proximal TFBSs with the gene, while motifs identified in the distal regulatory region of a gene are defined as distal TFBSs with the gene.

### Statistics and data visualization

If not specified, R (https://cran.r-project.org/;version 4.3.2) was used to compute statistics and generate plots if not specified. The Integrative Genomics Viewer (IGV) was used for the visual exploration of genomic data (Fig. 3g, 4b, 6g and 6i). The correlation between two groups of data was conducted with the Pearson analysis and *P* values are determined by the two-sided Pearson correlation coefficient analysis (Fig. 1e, 2h, 3e, Fig. S3b, 7b, 8b and 10e). For enrichment analysis, Fisher’s exact test was used, such as Fig. 2d, 5d, Fig. S5a, 10c. For the two groups’ comparison of data, the Student’s *t*-test was used such as Fig. 2, 3, 4, 5, 6, S4, Fig. S7, 8 and 9. For three or more independent groups comparison of data, Fisher’s Least Significant Difference was used in Fig. 2f and Fig. S4d.

**Fig. 1.**
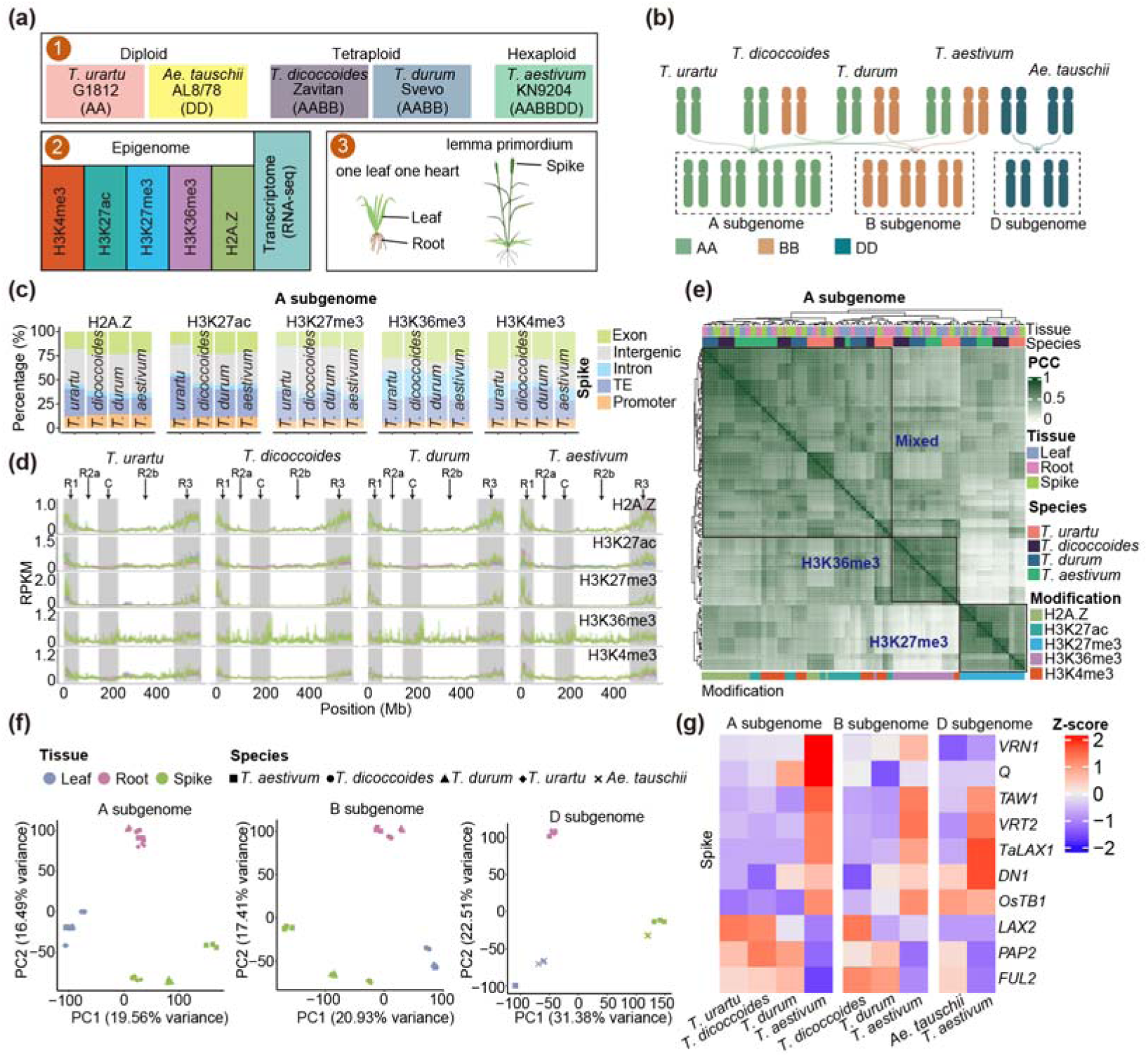
The epigenome and transcriptome profiles of hexaploid wheat and its tetraploid and diploid relatives. (a) Experimental design and the data series sampling across leaf, root and spike of hexa-, tetra- and dipoid wheat. (b) Strategy to integrate data for comparative analysis by splitting all species at the subgenome level. (c) Frequencies of histone mark peaks regions in diverse gene structural regions in A subgenome. The promoter is defined as distance TSS from -3,000 to 1,000bp. (d) Signals of histone markers along chromosome 1A in bins of 1Mbp length. R1, R2a, C, R2b, and R3 chromosomal regions are indicated within background colors. (e) Pearson’s correlation heatmap for the complete set of five histone modifications of A subgenome across all species and tissues. (f) Principal component analysis (PCA) of all expressed genes (TPM > 1) showing for different tissues and species. Each dot represents a sample. Three biological replicates were sequenced for each stage (g) Heatmaps showing normalized expression levels (TPM) of several known spike development related genes.

**Fig. 2.**
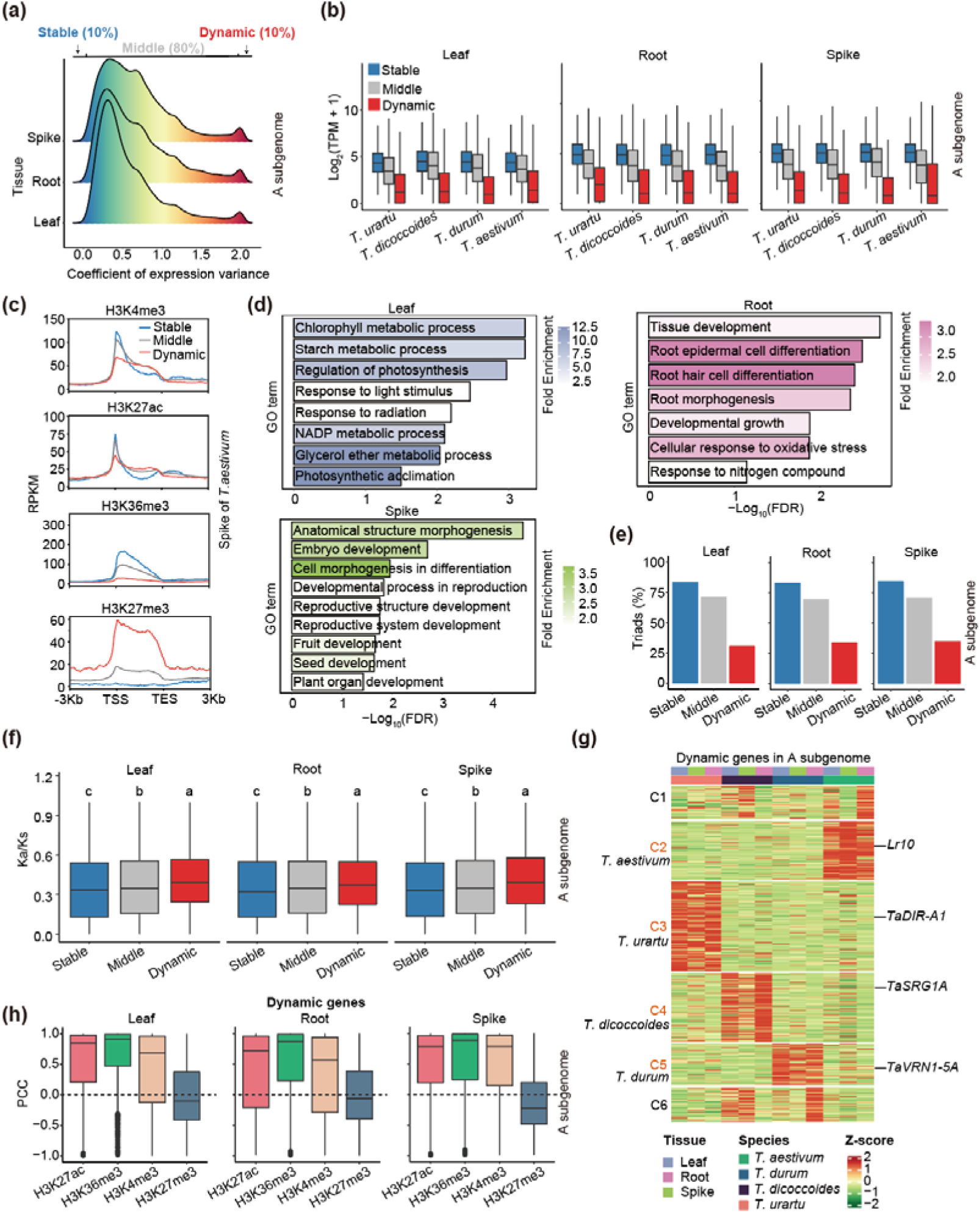
Characteristics of stable- and dynamic-expressed genes and associations with epigenetic modifications. (a) Classification of gene expression patterns in A subgenome based on coefficient of expression variations across all tissues and species. (b) Expression levels of stable, middle and dynamic genes across leaf, root and spike in A subgenome. (c) Histone modification landscape across stable, middle and dynamic genes. TSS, transcription starting site. TES, transcription ending site. (d) GO enrichment for stable genes across leaf, root, and spike in A subgenome. False discovery rate (FDR) by Benjamini-Hochberg procedure. (e) Percentages of triads genes in stable, middle, and dynamic genes in A subgenome. (f) Box plots of Ka/Ks ratios for stable, middle, and dynamic genes in A subgenome. The least significant difference (LSD) multiple comparison test was used to assess significance. Letters above the box denote significant differences (*P*≤0.05, ANOVA test). (g) Heatmap shown the expression levels of dynamic genes. Expression levels are normalized by z-scores of log_2_(TPM+ 1). Genes are grouped in clusters by *k*-means in A subgenome. Four species-specific expressed gene group are labelled in red, with represented genes notated. (h) Pearson’s correlation coefficients of four histone modifications across gene body of dynamic genes body between species with gene expression levels. The 3Kb flanking regions of gene bodies are included.

**Fig. 3.**
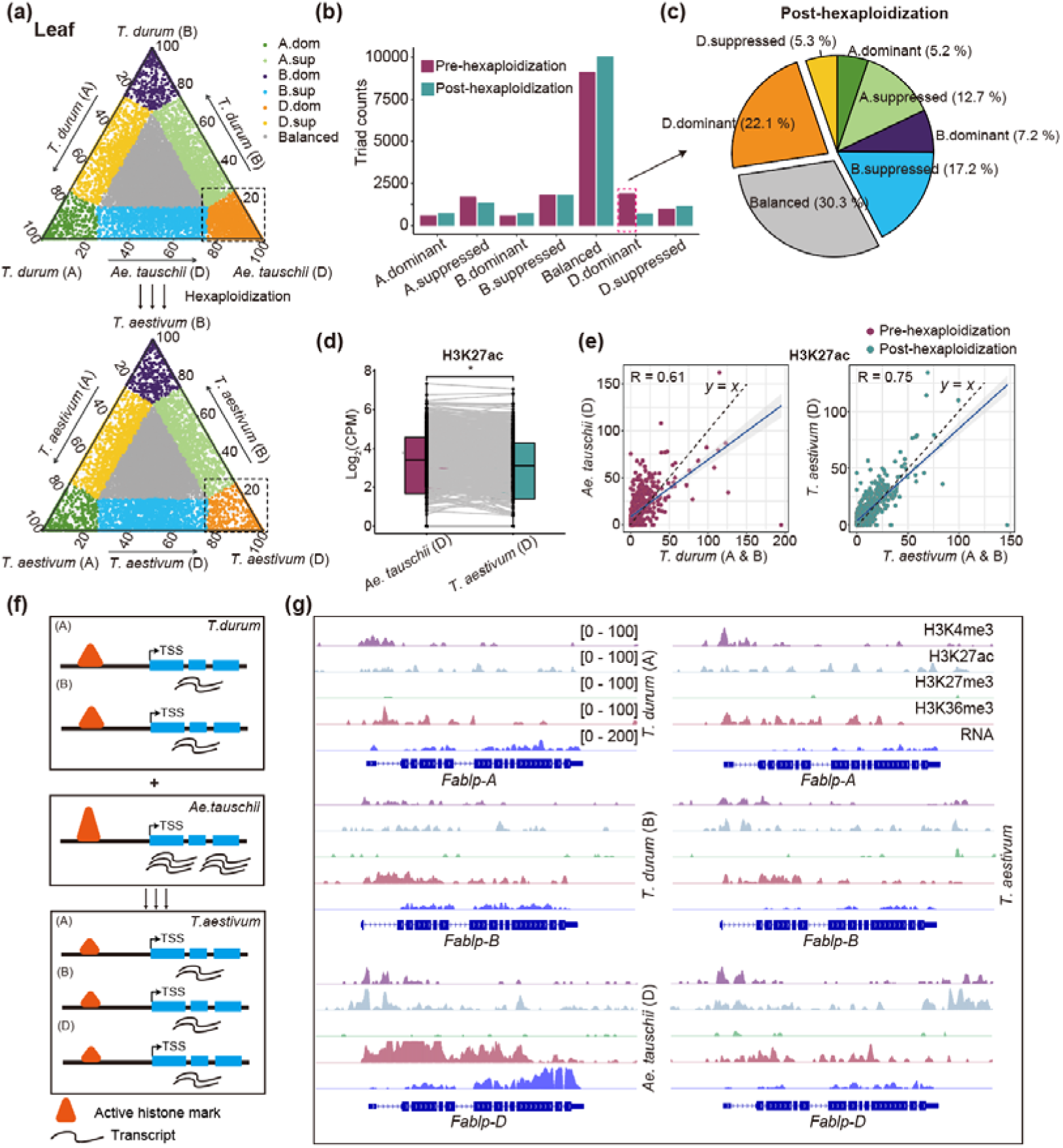
Repressed expression of genes newly introduced in wheat D subgenome. (a) Distribution of expression of orthologous genes between *T. durum* and *Ae. tauschii* (up) and homoeologous genes in A, B, and D subgenomes of *T. aestivum* (down) in leaf. Each circle represents an ortholog/homoeolog triad (composed of A, B, and D genome copies) coordinate consisting of the relative contribution of each homoeolog to the overall triad expression. (b) Counts of triads in seven categories pre- and post-hexaploidization. (c) Classification and percentage of triads in post-hexaploidization that were classified into D.dominant category in pre-hexaploidization. (d) Comparison of H3K27ac signal level of orthologs between *Ae. tauschii* and D subgenome in *T. aestivum*, which were classified into D.dominant to balanced categories before and after hexaploidization. Statistical significance was assessed by Student’s *t*-test. *, *P* ≤ 0.05. (e) Correlation of H3K27ac signals of orthologs between *Ae. tauschii* and *T. durum* (left) and homoeologs between D and A & B subgenomes in *T. aestivum* (right). The blue line denotes the line of best fit. (f) Schematic model of repressed- and convergent-expression of genes in the newly introduced in wheat D genome. (g) Genomic tracks illustrating the repression and convergence of expression and histone modification of *Fablp* genes in *Ae. tauschii, T. durum,* and *T. aestivum*.

**Fig. 4.**
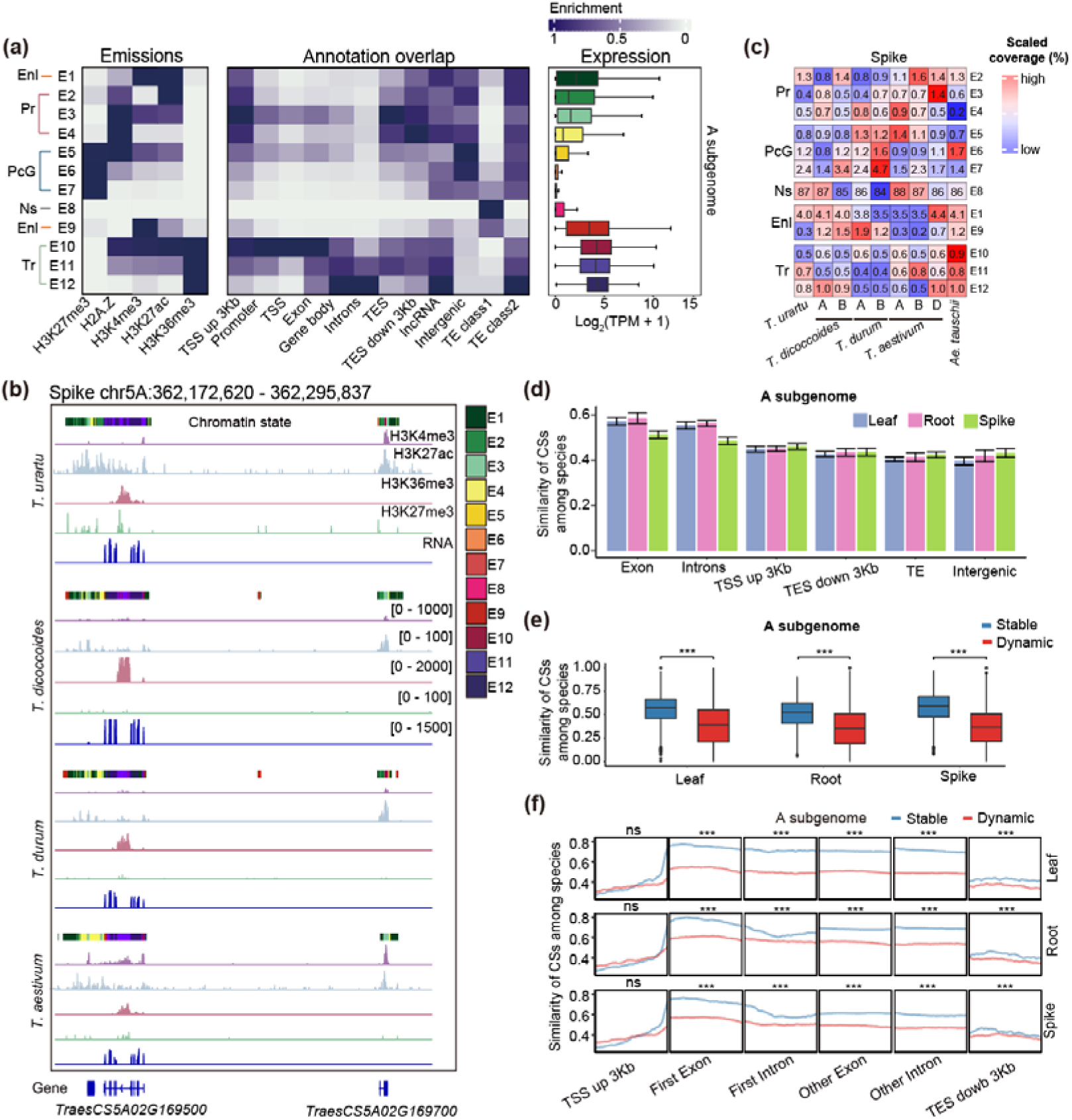
Chromatin states dynamic during the speciation of bread wheat. (a) Definitions and overview of chromatin state annotations (CSs) and epigenome data in A subgenome, which could be mainly categorized into five groups: Enl (enhancer-like), Pr (promoter), Tr (transcription), PcG (polycomb group), and Ns (no signals) state. Representative 12-chromatin state model based on five histone modification marks, emission probabilities for individual histone marks, and fold enrichments of chromatin states for the various types of genomic annotations and gene expression. (b) An example of chromatin state annotations. State colors are as shown in (A). (c) Comparison of the proportions of different CSs between the A, B and D subgenomes across five species in spike. (d) Similarity of CSs among species in different genomic regions across leaf, root, and spike in A subgenome. (e) Comparison of the similarity of CSs between stable and dynamic genes across leaf, root, and spike in A subgenome. Statistical significance was assessed by Student’s *t*-test. ***, *P* ≤ 0.001. (f) Distribution of similarity of CSs across gene body and flanking regions between stable and dynamic genes grouped by different transcription in A subgenome. Statistical significance was assessed by Student’s *t*-test. ***, *P* ≤ 0.001.

**Fig. 5.**
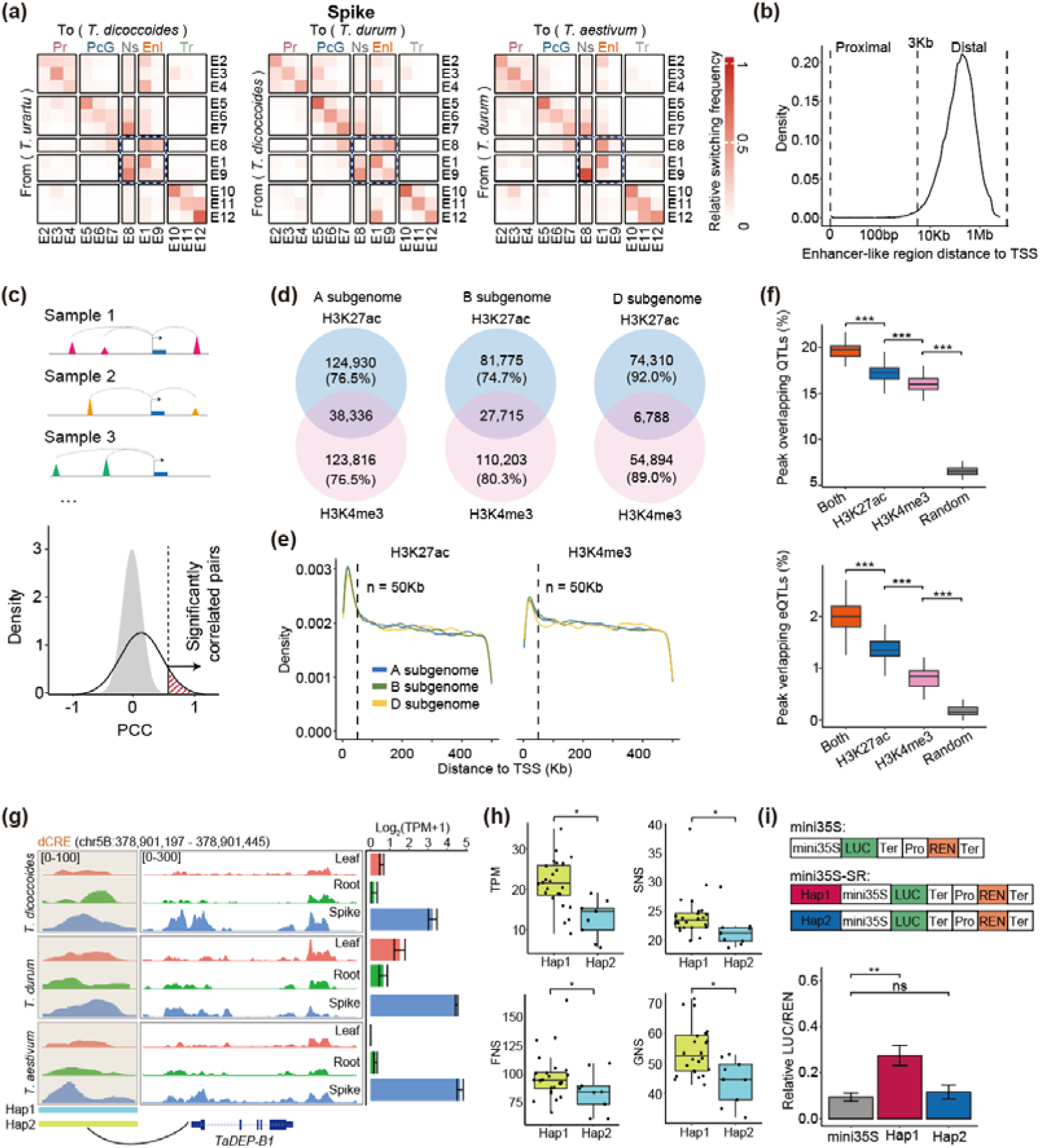
Association of traits with distal CREs mediated by H3K27ac and H3K4me3. (a) The switches of chromatin state during the polyploidization and evolution of wheat (from *T.urartu* to *T. dicoccoides*, *T. dicocccoides* to *T. durum*, and *T. durum to T. aestivum*). The color scale indicates the switching frequency. Ratio of the observed probability that a region switches from one CS (row) to another (column), The transition from Ns to Ns was removed. (b) Distribution of relative distance between the Enl and TSS. The x-axis is transformed to log10 scale. Regions within and over 3kb of the TSS were defined as proximal and distal, respectively. (c) Schematic overview of the computational strategy used to identify dCREs that are positively correlated with transcription of target genes. (d) Overlap of H3K27ac and H3K4me3-mediated dCRE identified in A, B, and D subgenomes. (e) Distribution of relative distance between the H3K27ac and H3K4me3-mediated dCRE and targeted genes. (f) Comparison of the peak percentage among H3K27ac-, H3K4me3-, H3K27ac-H3K4me3-co mediated dCREs, and random regions which were overlapped with QTLs and eQTLs. Statistical significance was assessed by Student’s *t*-test. ***, *P* ≤ 0.001. (g) The H3K27ac modification profile of *TaDEP-B1* with its distal dCRE. Expression data shown as mean ± s.d. of three biological replicates. (h) Comparison of the expression levels and spikelet number per spike (SNS), the number of florets per main spike (FNS) and the number of seeds per main spike (GNS) between two haplotypes in dCRE of *TaDEP-B1*. Statistical significance was assessed by Student’s *t*-test. ***, *P* ≤ 0.001. (i) Luciferase reporter assays validation of transcriptional regulatory role of distal region (Hap1 and Hap2) to *TaDEP-B1*. Statistical significance was assessed by Student’s *t*-test. ***, *P* ≤ 0.001.

**Fig. 6.**
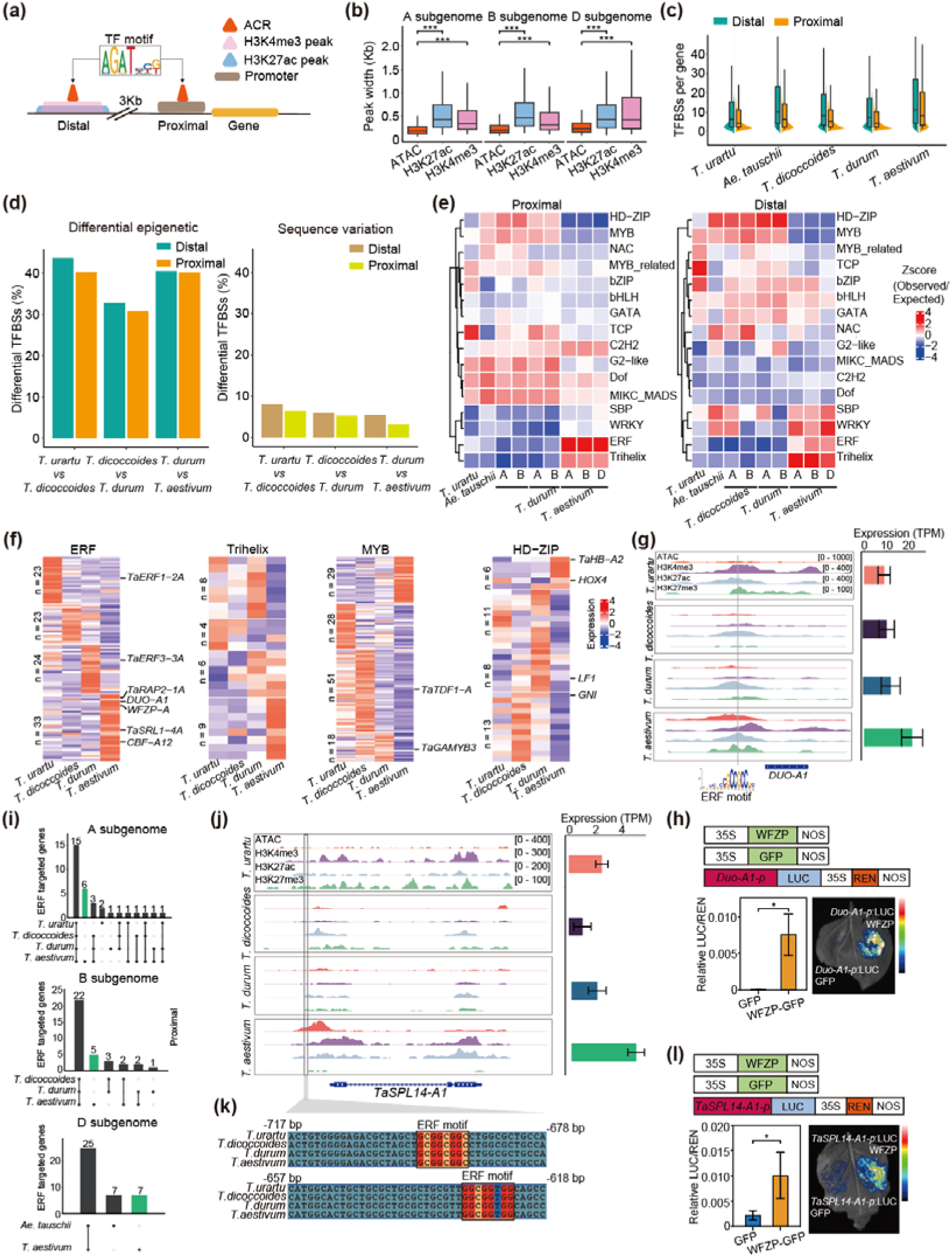
Changes of transcriptional regulatory relationships during the formation of hexaploid wheat. (a) Models of proximal and distal transcription factor regulatory relationship. (b) Comparison of the peak widths of ATAC, H3K27ac, and H3K4me3. Statistical significance was assessed by Student’s *t*-test. ***, *P* ≤ 0.001. (c) Distribution of proximal and distal transcription factor regulatory relationship counts (TRRs) for each gene across five species. (d) Ratio of changed TF binding sites due to the alterations of epigenetic modifications or sequence variations across different stages in A subgenome. (e) Motif density of clustered motifs from PlantTFDB in A, B and D three subgenome across five species. (f) Relatively expression levels of TFs belong to ERF, Trihelix, MYB, HD-ZIP families sorted by *k*-means clustering across the four species of spike in A subgenome. Known gene were labeled. (g) Genome tracks illustrating the chromatin accessibility, histone modification profile, and expression levels on the *DUO-A1* gene body and promoter. (h) Luciferase reporter assays show the transcriptional activation capability of WFZP (ERF family) to *DUO-A1*. Statistical significance was assessed by Student’s *t*-test. *, *P* ≤ 0.05, **, *P* ≤ 0.01, ***, *P* ≤ 0.001. (i) Overlap of spike-related genes (proximal) targeted by ERF TFs between different species in A, B, and D subgenomes. The dots at the bottom represent the types of intersections among each species layer. (j) Genome tracks illustrating the chromatin accessibility, histone modification profile, and expression levels on the *TaSPL14-A1* gene body and promoter, and ERF binding sites are labeled. (k) Alignment of the *TaSPL14-A1* promoter sequences of the four species, with the ERF motif highlighted. (l) Luciferase reporter assays show the transcriptional activation capability of WFZP (ERF family) to *TaSPL14-A1* in four species. Statistical significance was assessed by Student’s *t*-test. *, *P* ≤ 0.05, **, *P* ≤ 0.01, ***, *P* ≤ 0.001.

## Results

### Comprehensive profiling of the transcriptome and epigenome in common wheat and its relatives

The speciation of common wheat (*Triticum aestivum*, AABBDD) is marked by complicated polyploidization and domestication events, which are accompanied by genome-wide changes in genome structure, gene expression and epigenetic modifications (Song & Chen, 2015; Dvorak *et al*., 2018; Concia *et al*., 2020; Zhao *et al*., 2021). To unravel the regulatory dynamics between transcriptome and epigenome during wheat polyploidization and domestication, we applied RNA-seq and CUT&Tag (Cleavage Under Targets and Tagmentation) for core histone modifications in leaves, spikes, and roots tissues across multiple wheat species: *T. urartu* (AA, *c.v.* G1812)*, Ae. tauschii* (DD, *c.v.* AL8/78)*, T. dicoccoides* (AABB, *c.v.* Zavitan)*, T. durum* (AABB, *c.v.* Svevo) and *T. aestivum* (AABBDD, *c.v.* KN9204) (**Figs. 1a, Fig. S1a and Table S1**). While different species exhibited similar histone modification patterns, variations in signal intensity were observed (**Fig. S1b**). Principal component analysis (PCA) highlighted the high reproducibility among biological replicates, with same types of histone modifications clustering together more frequently (**Fig. S2**). Given the diverse genomic compositions of these species, we merged the A, B, and D subgenomes from different species and conducted comparative analyses at the subgenome level (**Fig. 1b**).

We annotated peaks of various histone modifications on genomic features from integrated three tissues and examined their distribution. H3K36me3 peaks were more enriched in gene bodies (**Fig. 1c, Fig. S3a and Table S2**). In contrast, H2A.Z, H3K27me3 and H3K27ac were predominantly localized in the ends of chromosomes, consistent with typical chromatin features of wheat reported in previous studies (Consortium *et al*., 2018; Zhao *et al*., 2023) (**Fig. 1d**). Correlation analysis of histone modifications across species and tissues revealed that H3K27me3 and H3K36me3 were prominently distinguishable from other modifications and exhibit strong species-specificity (**Fig. 1e and Fig. S3b**). For transcriptome, PCA revealed high similarity among samples from the same tissue, with greater variations observed in spike samples (**Fig. 1f**). For instance, key genes involved in spike development exhibited differential expression, such as *VERNALIZATION1* (*VRN1*) (Yan *et al*., 2003), *TAWAWA1* (*TAW1*) (Yoshida *et al*., 2013), *VEGETATIVE TO REPRODUCTIVE TRANSITION 2* (*VRT2*) (Kane *et al*., 2005), *FRUITFULL 2* (*FUL2*) (Li, C *et al*., 2019) and *Q* (Simons *et al*., 2006) (**Fig. 1g**).

Thus, we have generated comprehensive epigenomic and transcriptomic data across bread wheat and its relatives, enabling analysis of how epigenetic modifications drive transcription dynamics during speciation.

### Transcriptomic and epigenetic dynamics during common wheat speciation

To investigate transcriptomic dynamics during bread wheat speciation, we calculated the coefficient of expression variations (CEV) for genes within each tissue across species, classifying them into stable (lowest 10% CEV), dynamic (highest 10% CEV), and middle genes (remaining 80%) (**Fig. 2a, Fig. S4a and Table S3**). Stable genes exhibited higher expression levels than dynamic genes, consistent across the A, B, and D subgenomes (**Fig. 2b and Fig. S4b**). In spikes, active histone marks H3K4me3, H3K27ac, and H3K36me3 were enriched in stable genes, while H3K27me3 was depleted, indicating distinct epigenetic landscapes for gene groups (**Fig. 2c**).

Stable genes were enriched in tissue-specific Gene Ontology (GO) terms: photosynthesis and light response in leaves, root development and nitrogen response in roots, and reproductive growth and flower development in spikes (**Fig. 2d**), emphasizing their roles in tissue differentiation. Most stable genes were triads (containing A, B, and D subgenomes counterparts in hexaploid wheat (Ramírez-González et al., 2018)) (leaf: 83.5%, root: 83.1%, spike: 84.7%), while only ∼33% of dynamic genes were triads (**Fig. 2e and Fig. S4c**). Comparative non-synonymous (Ka) and synonymous (Ks) substitution ration (Ka/Ks) analysis revealed that stable genes are under stronger selective pressure, while dynamic genes experience relaxed selection (**Fig. 2f and Fig. S4d**). For instance, the leaf senescence gene *ascorbate peroxidase 4* (*TaAPX4-A1*) (Ribeiro *et al*., 2017) in leaf, the nitrate transporter gene *chloride channel protein 2* (*TaKNCLC-2A*) (Shi *et al*., 2022) in roots, and the spike development gene *squamosa promoter binding protein like 15* (*TaSPL15-A*) (Pei *et al*., 2022) maintains stable expression, all of which endure significant selective pressures (**Fig. S4e**).

Dynamic genes showed species-specific expression patterns. Using K-means clustering, we identified groups with across tissues expression in specific species such as *T. aestivum* (C2), *T. urartu* (C3), *T. dicoccoides* (C4), and *T. durum* (C5) (**Fig. 2g**). GO analysis highlights the enrichment of pathways related to biotic stress-related pathways in *T. aestivum*, such as the rust resistance gene *leaf rust 10* (*Lr10*) (Feuillet *et al*., 2003). In *T. durum*, pathways associated with inorganic salt metabolism and stress responses were enriched, including the *cadmium ion tolerance 1* (*TaCDT1*) (Kuramata *et al*., 2009) (**Fig. S5a**). *T. dicoccoides* exhibited enrichment of pathways related to cellular aldehyde metabolism, such as the *alcohol dehydrogenase 1* (*TaAdh1*) (Wang, S *et al*., 2021). *T. urartu* showed enrichment of pathways related to abiotic stimulus and temperature stress, including the *heat shock protein 18.2* (*TaHSP18.2*) (Kaur *et al*., 2015). Such pattern was consistent in subgenomes B and D (**Fig. S5b**), with *Ae. tauschii* showing the highest number of species-specific genes (**Fig. S5c**). Histone modifications strongly correlated with expression dynamics. Active mark (H3K27ac, H3K36me3, H3K4me3) positively correlated with dynamic gene expression change across species, while the repressive mark H3K27me3 showed a negative correlation (**Fig. 2h**).

Thus, stable genes during bread wheat speciation exhibit conserved coding sequences and subgenome triads, maintaining tissue identity. In contrast, dynamic genes reflect species-specific biological processes, with active histone modifications driving their expression, underscoring the significance of chromatin landscapes in wheat evolution.

### Epigenetic regulation balances subgenome expression during common wheat hexaploidization

To understand the impact of polyploidization on homoeolog/ortholog expression, we analyzed triads before and after hexaploidization by combining the orthologous of *T. durum* (AABB) and *Ae. tauschii* (DD) to form pre-hexaploidization triads, and the A, B, and D subgenomes of *T. aestivum* (AABBDD, *c.v.* KN9204) for post-hexaploidization triads. Using 16,510 expressed triads in leaf tissue (Ramírez-González et al., 2018), we categorized them into seven groups: one balanced group (similar expression across homoeologs/orthologous) and six dominant or suppressed groups based on relative expression levels of individual homoeolog/orthologous (**Fig. 3a**).

Although most triads exhibited balanced expression patterns both pre- and post- hexaploidization, the number of balanced triads increased from 9,106 to 10,055 post-hexaploidization. Concurrently, D subgenome-dominant triads decreased significantly, from 1,816 to 700 (**Fig. 3b and Table S4**). Notably, 30.3% of D - dominant triads pre-hexaploidization shifted to balanced expression post-hexaploidization, while 22.1% remained D-dominant (**Fig. 3c**). Similar reductions in D-dominant triads were observed in roots but were less pronounced in spikes, possibly due to the extended developmental stages of spikes and more complex transcriptional regulation (**Fig. S6a, 6b and Table S4**). These findings suggest a repression of D subgenome homoeologs/orthologs were during hexaploidization to maintain expression balance across subgenomes.

To elucidate the mechanisms behind this shift, we examined epigenomic changes in leaves. Post-hexaploidization, D-dominant-to-balanced (Dd-to-Db) triads exhibited decreased levels of active histone marks, including H3K27ac, H3K36me3, and H3K4me3, while inhibitory H3K27me3 levels remained unchanged (**Fig. 3d** and **Fig. S7a**). By contrast, D-dominant-to-D-dominant (Dd-to-Dd) triads showed no significant changes in histone modifications (**Fig. S8a**). Additionally, correlations in H3K27ac levels between the D and A/B subgenomes increased from 0.61 to 0.75 during hexaploidization (**Fig. 3e**), a trend also observed for H3K4me3, H3K36me3, and H3K27me3 (**Fig. S7b**). And this trend was not observed in Dd-to-Dd triads (**Fig. S8b**).

We propose that during hexaploidization, the fusion of the D subgenome from *Ae. tauschii* with tetraploid wheat reduced the necessity for high expression levels characteristic of the diploid state. Increased gene copy numbers allowed for convergence of expression levels and epigenomic profiles across subgenomes (**Fig. 3f**). For example, genes such as *Falbp* (Bak *et al*., 2013), involved in stomatal closure, exhibited higher expression in *Ae. tauschii* but aligned with A and B subgenome expression levels post-hexaploidization (**Fig. 3g**).

In conclusion, bread wheat hexaplodization rebalanced gene expression among subgenomes by reducing the levels of active histone marks in the D subgenome. This convergence highlights the adaptive mechanisms underlying gene regulation during polyploidization, ensuring the stability and functionality of the hexaploid genome.

### Chromatin state similarity aligns with transcriptional stability during common wheat speciation

Given that histone modifications often work together to regulate transcription, we further analyzed chromatin states (CSs) by integrating combinatorial patterns of multiple histone marks to annotate the epigenome and characterize transcriptional activity (Lee *et al*., 2010; Ernst & Kellis, 2012; Sequeira-Mendes *et al*., 2014). Using ChromHMM, we trained a 12-CS model at a 200-bp resolution for three tissues across all species, incorporating profiles of four core histone modifications and the H2A.Z variant. The identified CSs (E1-E12) were further grouped into five major functional categories: promoter (Pr), enhancer-like (EnL), transcriptional (Tr), polycomb group (PcG), and no signal (Ns) (**Fig. 4a and Fig. S9a**). For instance, E2, E3, and E4, marked by H2A.Z and H3K27ac at transcription start sites (TSS), were classified as Pr. While E1 and E9, enriched for H3K4me3 and H3K27ac, were categorized as EnL. PcG states (E5, E6, and E7) were marked by H3K27me3, and Tr states (E10, E11, E12) were defined by H3K36me3 and associated with high-expression genes. E8, lacing modification signals, was categorized as Ns. Across subgenomes A, B, and D, the genomic characteristics and expression patterns of each chromatin state were largely conserved, indicating a shared epigenetic code (**Fig. 4a**, **Fig. 4b and Fig. S9a**).

Although the overall distribution of CSs across the five species were consistent, sub genome-specific variations were observed (**Fig. 4c, Fig. S9b and Table S5**). Notably, the Enl and PcG states in the D subgenome of *T. aestivum* more closely resembled those of *Ae. tauschii* than the A and B subgenomes of *T. aestivum* (**Fig. 4c and Fig. S9b**), likely due to the shorter evolutionary history of the D subgenome. Genome-wide comparisons revealed that CSs around gene bodies and their flanking regions were more similar between hexaploidy wheat and its tetraploid and diploid relatives than TE and intergenic regions (**Fig. 4d and Fig. S9c**).

Stable genes exhibited the highest conservation of CSs across species in leaves, roots, and spikes, whereas dynamic genes showed greater variability (**Fig. 4e and Fig. S9d**). To explore this further, gene regions and their flanking sequences were divided into six segments: upstream 3kb of the TSS, the first exon, the first intron, other exons, other introns, and downstream 3kb of the transcription end site (TES). Differences in CSs similarity between stable and dynamic genes were most pronounced within the gene body regions, while upstream and downstream regions showed smaller or nonsignificant difference. These findings underscore the critical role of chromatin states within gene bodies in driving expression changes during bread wheat speciation (**Fig. 4f and Fig. S9e**).

### Distal region chromatin state dynamics associate with agronomic trait variations

To explore chromatin state dynamics during bread wheat speciation, we analyzed transitions between CSs across species in different tissues. In the A subgenome, frequent transitions between Ns (E8) and Enl (E1 and E9) states were observed across leaves, roots, and spikes during the progression from *T. urartu* to *T. dicoccoides*, *T. dicoccoides* to *T. durum*, and *T. durum* to *T. aestivum* (**Fig. 5a, Fig. S10a and Table S6**). Enl states, often located over 3 kb from genes, likely functions as distal regulatory elements (**Fig. 5b**). To investigate their role in gene expression regulation during wheat evolution, we examined the correlation between distal H3K27ac/H3K4me3 peak (located between 3-500 kb from gene loci) and gene expression across all species. Gene-peak pairs with significant correlations (*P<*0.05) (Corces *et al*., 2018) were identified as candidate distal regulatory relationships (**Fig. 5c**).

We identified 388,228 H3K27ac-mediated gene-distal *cis* regulatory elements (dCREs) pairs (A: 163,265, B: 109,489, D: 80,781) and 360,683 H3K4me3-mediated pairs (A: 161,760, B: 137,239, D: 61,681), involving 43,431 to 100,757 histone peaks and 15,366 to 19,250 genes (**Fig. S10b and Table S7**). Approximately 10∼25% of dCRE pairs overlapped between H3K27ac and H3K4me3 modifications (**Fig. 5d**). dCREs were enriched within 50 kb of gene loci, with H3K27ac-mediated interactions tending to be shorter than those mediated by H3K4me3 (**Fig. 5e**).

To understand the functional significance of these dCREs, we compared them with quantitative trait locus (QTL) and expression QTL (eQTL) regions associated with agronomic traits. dCREs co-mediated by H3K27ac and H3K4me3 were most significantly enriched in these regions (**Fig. 5f**), suggesting their pivotal role in regulating key traits. GO analysis indicated that genes regulated by both H3K27ac and H3K4me3 marks were enriched in pathways related to biotic and abiotic stress and organ development (**Fig. S10c**). *T*. *aestivum* exhibited the highest number of species-specific dCREs (**Fig. S10d**), likely due to regulatory integration from three subgenomes following hexaploidization. These species-specific dCREs showed higher frequencies of species-specific single nucleotide polymorphisms (SNPs), suggesting that sequence variation underpins differences in dCREs across species (**Fig. S5e**).

For instance, we identified a dCRE significantly associated with the expression of *TaDEP-B1* (Huang *et al*., 2009) (**Fig. 5g and Supplementary Fig. 10f**), which influences grain number per spike. In hexaploid wheat populations, this dCRE exists in two haplotypes (Hao *et al*., 2020; Zhou *et al*., 2020) (**Supplementary Fig. 10g**). Hap1 exhibited significantly higher *TaDEP-B1* expression, spikelet number per main spike (SNS), florets per spike (FNS), and grains per spike (GNS) compared to Hap2 (Wang, Y *et al*., 2017) (**Fig. 5h**). Luciferase reporter assays further confirmed the enhancer like role of Hap1’s dCRE, while Hap2 lacked this function (**Fig. 5i**).

In conclusion, H3K27ac- and H3K4me3-mediated modifications in distal regulatory regions play a crucial role in trait variation.

### Expansion of ERF transcriptional regulation in common wheat is involved in shaping spike traits

Epigenetic modifications often couple with transcription factors (TFs) activity dynamics, where accessible chromatin regions (ACRs) serve as docking sites for TFs regulating gene transcription (Davie *et al*., 2015; Schmitz *et al*., 2021) (**Fig. 6a**). To explore TF-mediated regulatory changes, we generated ATAC-seq data for hexaploid wheat and its relatives, focusing on the spikes due to their significant morphological evolution. We identified 75,635 gene body ACRs (gACRs), 134,385 promoter ACRs (pACRs), and 349,537 distal ACRs (dACRs), with more than half located distally (**Fig. S11a**). ATAC peaks were narrower than those for H3K4me3 and H3K27ac, offering enhanced resolution for detecting TF binding sites (**Fig. 6b**). pACRs were classified as proximal open regulatory regions, while dACRs within H3K27ac- or H3K4me3-mediated dCREs were designated as distal open regulatory regions (**Fig. 6a**).

A genome-wide scan for TF binding motifs identified transcription factor binding sites (TFBSs) within open regulatory regions across five species. Distal TFBSs outnumbered proximal ones, and hexaploid wheat exhibited the most TFBSs, likely reflecting its integrated regulatory networks from three subgenomes (**Fig. 6c**). To understand how chromatin modifications and sequence variations shaped TFBSs during bread wheat speciation, we distinguished TFBS changes caused by sequence variations versus epigenetic modifications (e.g., ACR, H3K27ac, or H3K4me3) (**Fig. S11b**). Epigenetic-driven TFBSs changes accounted for 30-50% of the differences among species, while sequence-driven changes were less than 10%, underscoring the pivotal role of epigenomic landscape dynamics in TFBS evolution (**Fig. 6d, Fig. S11c and 11d**).

Examining TFBS density across proximal and distal regulatory regions revealed consistent patterns within subgenomes of the same species, reflecting balanced transcriptional regulation among subgenomes. However, significant differences were observed between *T. aestivum* and its diploid and tetraploid relatives (**Fig. 6e**). Notably, *T. aestivum* gained 14,019 and 2,002 TFBSs for the ethylene-responsive factor (ERF) and Trihelix TF families in proximal regions and 14,019 and 2,002 in distal regions, while losing 9,285 and 921 TFBS for the MYB and HD-ZIP families in proximal regions and 9,314 and 1,325 in distal regions (**Fig. S11e**). This suggests a unique role for the ERF family in spike development in hexaploid wheat.

To investigate whether TFBS specificity in hexaploid wheat aligns with TF expression, we analyzed four TF families. Hexaploid wheat showed elevated expression of ERF and Trihelix TFs, including spike regulators such as DUO and FRIZZY PANICLE (WFZP) (Du *et al*., 2021; Wang *et al*., 2022), alongside reduced MYB and HD-ZIP TFs expression (**Fig. 6f**). This synchrony between TFBS expansion and TF expression levels suggests a coordinated regulatory evolution, a pattern consistent in the B and D subgenomes (**Fig. S12a and 12b**). For instance, the specific expression of *DUO-A1* in hexaploid wheat associated with enhanced chromatin accessibility, increased active histone modification signals (**Fig. 6g**). Interestingly, an ERF binding site was identified within the *DUO-A1*’ opened promoter region. Luciferase assays demonstrated the transcriptional activation of WFZP to *DUO-A1* promoter, suggesting the presence of a complex regulatory circuit within the ERF family (**Fig. 6h**).

Further, analysis of the 283 high-confidence spike development-related genes (Luo *et al*., 2023; Lin *et al*., 2024) revealed ERF binding sites in the proximal region of 107 genes and in the distal regions of 108 genes, with 18 proximal and 24 distal sites exclusively present in *T. aestivum* (**Fig. 6i, Fig. S12c and Table S8**). For example, increased chromatin accessibility in the promoter of *TaSPL14-A1* in *T. aestivum* covered two ERF binding sites, which may be linked to higher expression (**Fig. 6j**). Notably, these binding sites exist across species with no sequence variations (**Fig. 6k**), emphasizing chromatin accessibility as the key driver instead of the DNA sequence *per se* in this context. Functional validation via luciferase assays confirmed these findings (**Fig. 6l**).

These data reveal the co-evolution of TFs and their binding sites during wheat speciation, highlighting the expansion of ERF family binding sites within open chromatin regions of genes in *T. aestivum* underpinning the spike development.

## Discussion

### Epigenetic regulation of transcription during common wheat speciation

Epigenetic modifications are critical for regulating transcription, growth, and developmental processes in plants (Roy *et al*., 2010; Dunham *et al*., 2012). Here, we constructed a multi-tissue epigenomic and transcriptomic map of wheat’s A, B, and D subgenomes across diploid, tetraploid, and hexaploid species. Our findings reveal two types of gene expression pattern during common wheat speciation: stable and dynamic expression.

Genes with stable expression, such as the spike development gene *TaSPL15-A*, are enriched in conserved pathways linked to tissue-specific functions. In contrast, dynamically expressed genes, like the leaf rust resistance gene *Lr10*, which is specific to hexaploid wheat, are associated with stress responses and secondary metabolism, often driving individual fitness variations (Fig.2a). Stably expressed genes exhibit higher transcription levels, stronger activating histone marks (H3K4me3, H3K27ac, H3K36me3), lower repressive marks (H3K27me3), and more conserved amino acid sequences, while dynamic genes display the opposite trends (Fig.2b, 2c, and 2f). This suggests a coordinated evolution of gene expression, histone modification, and sequence conservation, promoting functional innovation. We also observed that dynamic gene expression differences across species align with histone modification patterns (Fig.2h, 4e, and 4f), highlighting epigenetic flexibility as a key driver of adaptability without altering DNA sequences (Jaenisch & Bird, 2003).

During the formation of hexaploid wheat, merging with the D subgenome led to widespread gene expression changes (Vasudevan *et al*., 2023) (Fig.3a-c). Activating histone marks of *Ae. tauschii* genes were suppressed, reducing their transcription levels and aligning them with homologous genes from the A and B subgenomes. This “diploidization” of hexaploid wheat may enhance genomic stability and heterosis. Differential epigenetic modifications play a crucial role in driving subgenome functional differentiation by shaping the environment for co-factor binding (Wang, M *et al*., 2021) and influencing chromatin interactions (Concia et al., 2020). These processes are further impacted by the integration of co-factors from fused subgenomes, highlighting the complexity of regulatory mechanisms. Our findings underscore the significance of epigenomic modifications and subgenomic interactions in wheat breeding. This knowledge provides valuable insights for selecting parental lines and designing genetic materials in chromosomal engineering to enhance crop traits and adaptability

### Distal enhancer-like regions regulate gene expression and phenotypic variation

Despite advancements in wheat epigenomics, our understanding of chromatin changes and their coordinated effects on gene expression during wheat polyploidization and domestication remains limited (Xiang *et al*., 2019; Yuan *et al*., 2020; Liu *et al*., 2021; Yuan *et al*., 2022; Miao *et al*., 2024). Here, we identify frequent transitions in distal chromatin regions enriched with H3K4me3 and H3K27ac marks (Fig.5a). These marks, associated with potential enhancers (Li, Z *et al*., 2019; Zhang *et al*., 2023; Zhao *et al*., 2023), suggest a flexible regulatory mechanism underpinning gene expression changes during wheat speciation.

Our study establishes a link between distal regulatory regions and gene expression, revealing significant overlaps between H3K27ac- and H3K4me3-marked regions with QTL and eQTLs (Fig.5f). These findings highlight the biological significance of distal elements in gene regulation. For instance, we discovered an H3K27ac-mediated regulatory site of the spike grain number regulator TaDEP-B1. Two haplotypes in modern wheat varieties lead to significant differences in *TaDEP-B1* expression and spikelet number (Fig.5g-i). This underscores the critical role of distal regulatory regions in shaping agronomic traits

While distal regulatory relationships identified through epigenetic signal correlation lack direct physical evidence, emerging 3D genomics technologies offer promising tools to refine long-range regulatory networks (Yuan *et al*., 2022; Deng *et al*., 2023), providing deeper insights into wheat genome regulation.

### The co-evolution of TFs and their binding sites during wheat speciation

Molecular evolution drives genetic differentiation, ontogeny, and trait persistence (Cui *et al*., 2021). The co-evolution of transcription factors and their binding sites plays a critical role in this process. Wheat speciation is accompanied by dynamic changes in transcriptional regulatory networks, as seen in previous co-expression studies of diploid, tetraploid, and hexaploid wheat (Li *et al*., 2023). However, these studies did not explore how TF binding depends on chromatin environments and matching binding sequences (Trojanowski & Rippe, 2022).

Acquiring new TFBSs through sequence variations poses a significant challenge for organisms (Tuğrul *et al*., 2015). Our research shows that epigenomic changes significantly impact TFBSs, far outweighing the effects of DNA sequence alterations (Fig.6d). Polyploidization introduces substantial epigenetic changes, and modulating TFBSs through chromatin modifications is more efficient, flexible, and cost-effective approach than DNA changes. In hexaploid wheat, we observed an increase in ERF and Trihelix TFBSs, alongside a decrease in MYB and HD-ZIP TFBSs (Fig.6e). Notably, corresponding changes in the expression levels of these TFs were observed during wheat speciation (Fig.6f). This reflects a potential co-evolutionary relationship between TFBSs and TFs during the evolutionary processes of both animals and plants (Zhang *et al*., 2022).

Previous studies have suggested a co-evolutionary link between TFs and TFBSs (Yang *et al*., 2011), but the relationship between the expansion and contraction of TFBSs and TF expression has not been explored. In the case of ERF TFs, increased expression corresponds with altered chromatin states at ERF binding sites, making them more accessible and thus activating gene expression (Fig.6g-l). This indicates intricate feedback mechanisms driving the co-evolution of TFs and TFBSs, contributing to the evolution of transcriptional regulatory networks during wheat speciation.

## Acknowledgements

We thank Prof. Hong-Qing Ling for the G1812 seeds, Prof. Aimin Zhang for the AL8/78 seeds, and Prof. Fei Lu for the Zavitan seeds, all from Institute of Genetics and Developmental Biology, Chinese Academy of Sciences. And we thank Prof. Shisheng Chen for the Svevo seeds from Peking University.

## Author contributions

J.X. designed and supervised the research, J.X., Z.-H. Z., X.-L, L. wrote the manuscript. X.-L. L. performed CUT&Tag, ATAC-seq and RNA-seq experiments; Z.- H, Z., and Y.-X. X., performed bioinformatics analysis; J.-J. Y. did transcription regulation assay. W.-Q. T., L.-F. M., W.-L. G. revised the manuscript; Z.-H. Z., and J.X. prepared all the figures. All authors discussed the results and commented on the manuscript.

## Data and code availability

Sequencing data of CUT&Tag, ATAC-seq and RNA-seq are available at Genome Sequence Archive of the National Genomics Data Center, China National Center (<u>PRJCA027414</u>) and are publicly accessible at https://ngdc.cncb.ac.cn/gsa. CUT&Tag, ATAC-seq and RNA-seq data for spike and root tissues of KN9204 were published previously (Lin et al., 2024; Zhang et al., 2023). Code used for all processing and analysis is available in GitHub (https://github.com/ZhangZhaoheng24/246-epigenome).

## Supplemental information

Supplemental Figures 1□12 and Supplemental Tables 1-9.

**Fig. S1.**
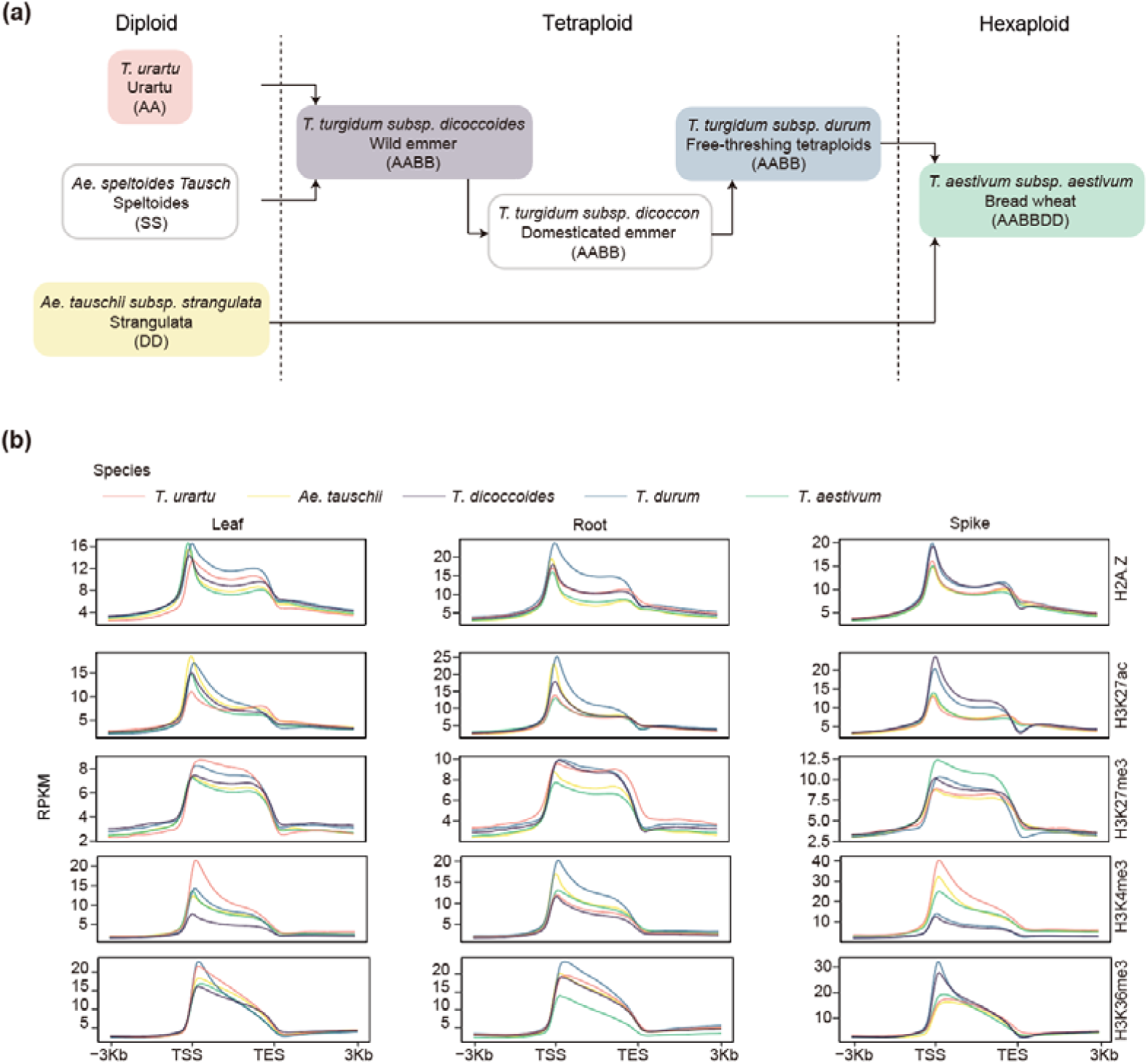
The species used in this study and their histone mark profiles in the gene body and flanking regions. **(a)** Schematic diagram of the speciation process of bread wheat through two successive rounds polyploidization. Where the species we used in this study are colored in. **(b)** Density of histone modifications in relation to the gene body as well as regions surrounding ± 3 Kb of the transcription start (TSS) and end (TES) sites.

**Fig. S2.**
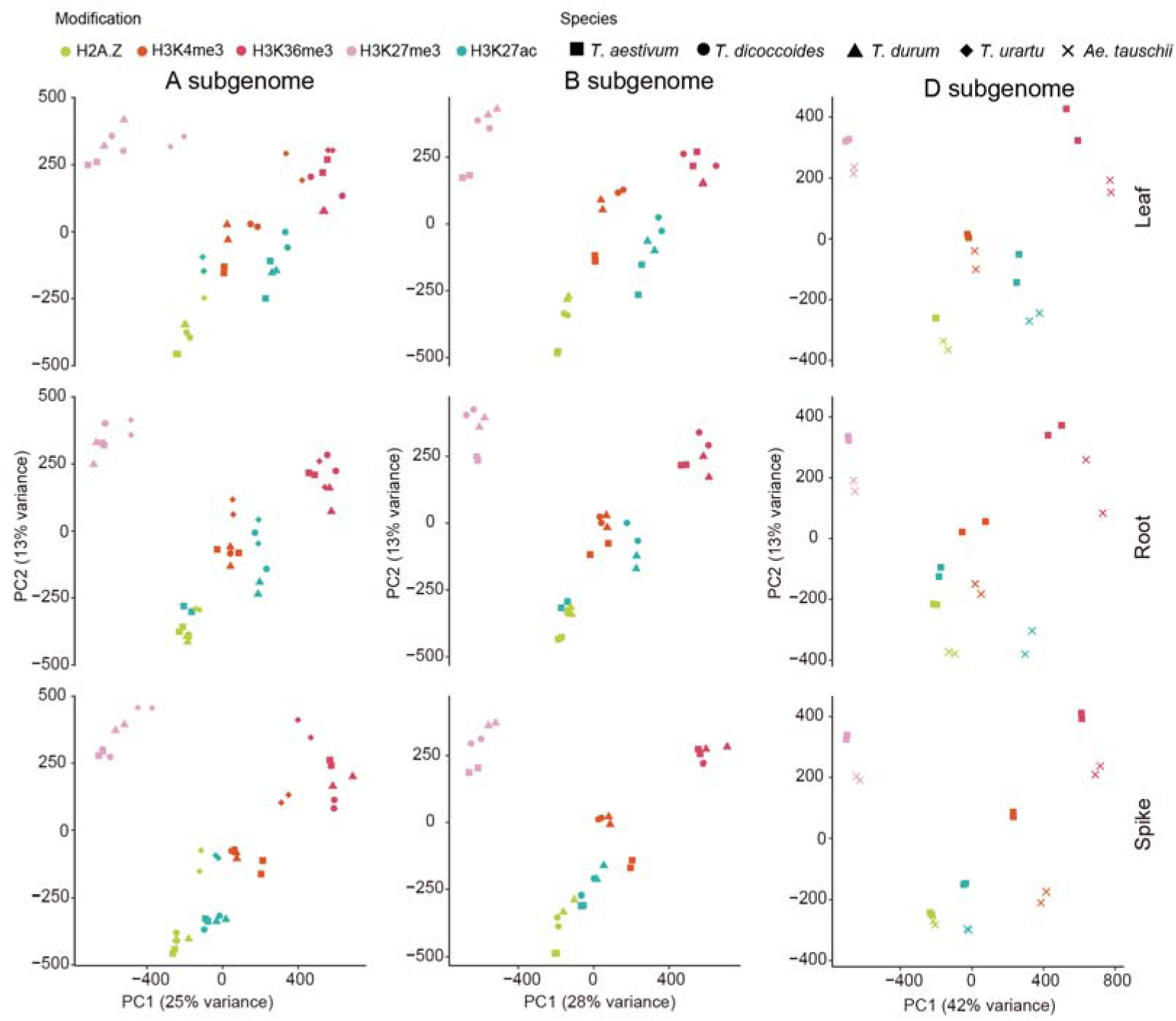
PCA of genome-wide epigenetic modification (H3K27ac, H2A.Z, H3K36me3, H3K27me3, H3K4me3) levels showing for different tissues and species. Each dot represents a sample. Three biological replicates were sequenced for each stage.

**Fig. S3.**
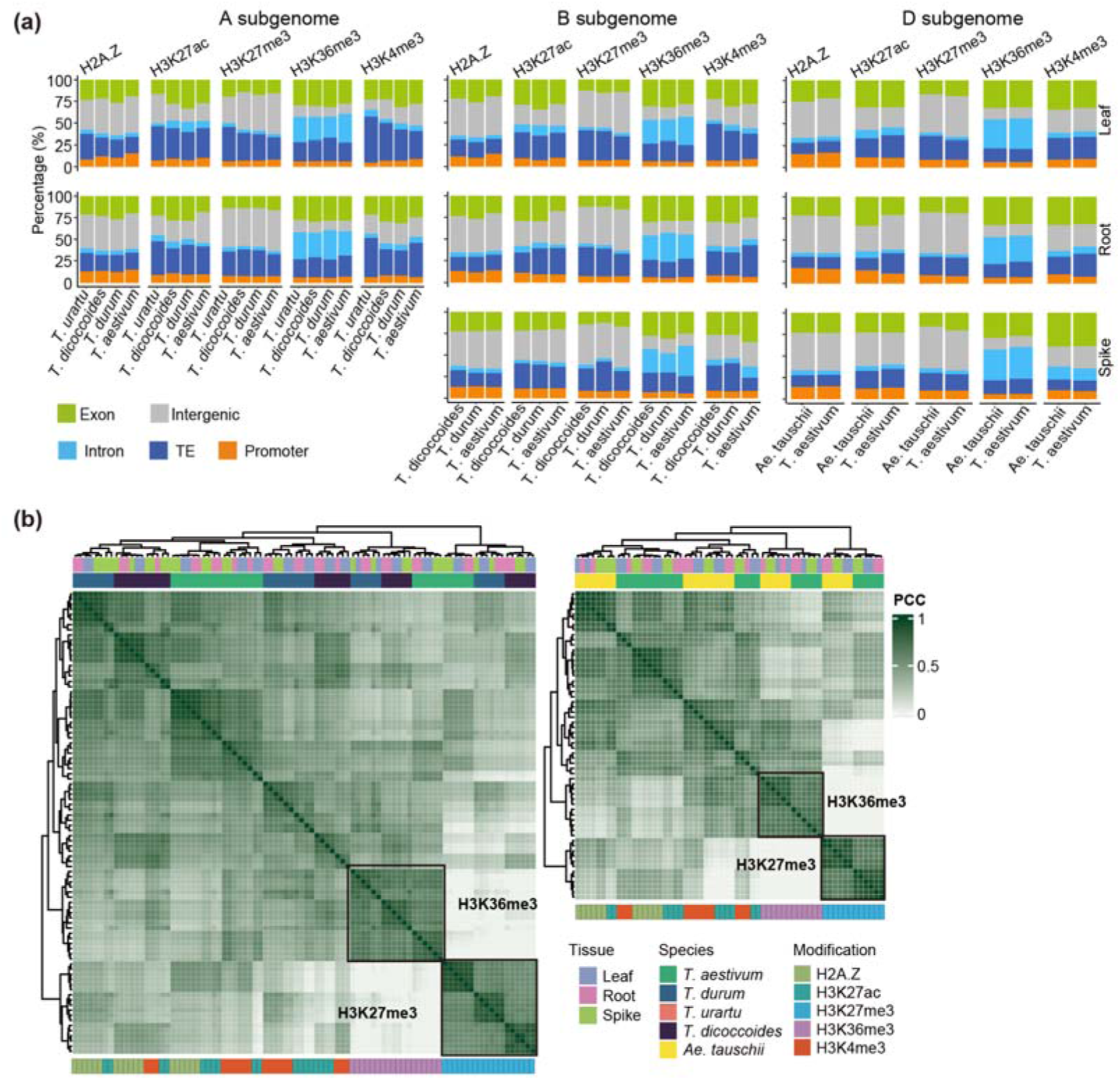
Peak distribution and correlation of histone marks on different genomic features. (a) Frequencies of histone mark peaks regions in diverse gene structural regions in B and D subgenome. The promoter is defined as distance TSS from -3,000 to 1,000bp. (b) Pearson’s correlation heatmap for the complete set of five histone modifications of B and D subgenome across all species and tissues.

**Fig. S4.**
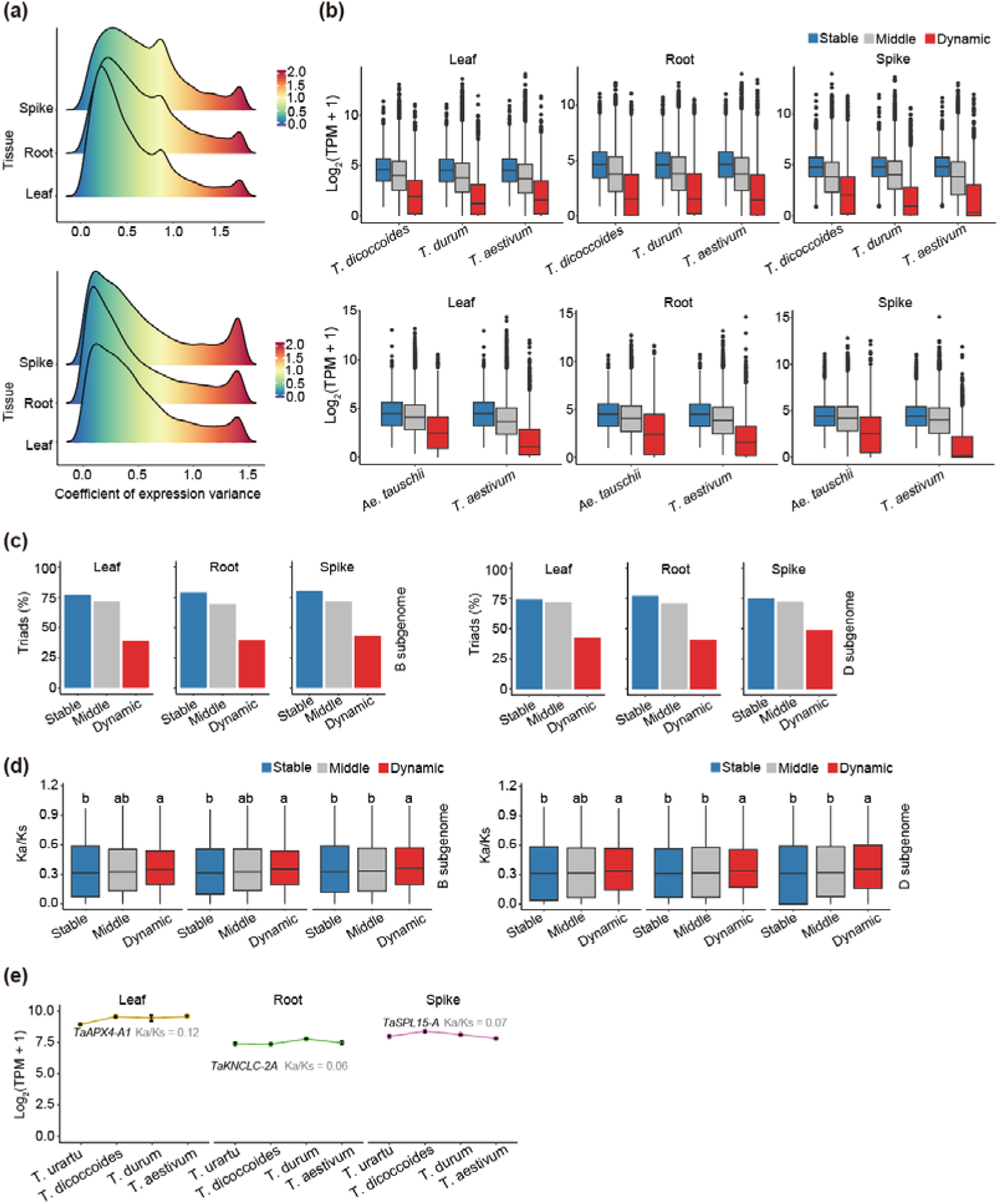
Characteristics of stable and dynamic genes in B and D subgenome. (a) Classification of gene expression patterns in B and D subgenome based on coefficient of expression variations across all tissues and species. (b) Expression levels of stable, middle and dynamic genes across leaf, root and spike in B and D subgenome. (c) Percentages of triads genes in stable, middle, and dynamic genes in B and D subgenome. (d) Box plots of Ka/Ks ratios for stable, middle, and dynamic genes in B and D subgenome. The least significant difference (LSD) multiple comparison test was used to assess significance. Letters above the box denote significant differences (*P*≤0.05, ANOVA test). (e) Gene expression and Ka/Ks of representative genes in leaf, root and spike.

**Fig. S5.**
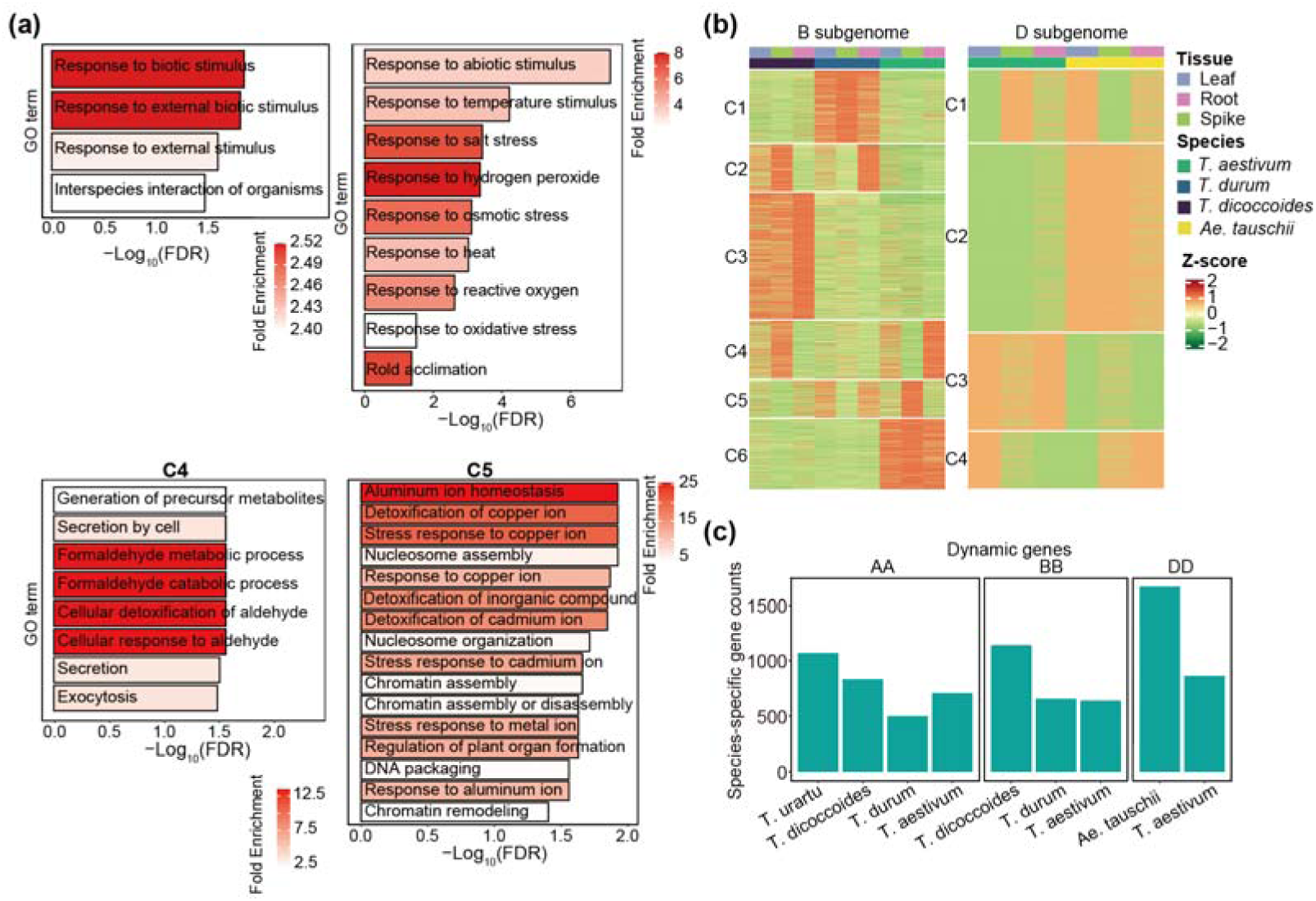
Characteristics of dynamically expressed genes. (a) GO enrichment for genes in C2, C3, C4 and C5 from Figure 2G. False discovery rate (FDR) by Benjamini-Hochberg procedure. (b) Heatmap shown the expression levels of dynamic genes. Expression levels are normalized by z-scores of log_2_(TPM+ 1). Genes are grouped in clusters by *k*-means in B and D subgenome. (c) The number of species-specific expressed genes across four speices and A, B and D three subgenomes.

**Fig. S6.**
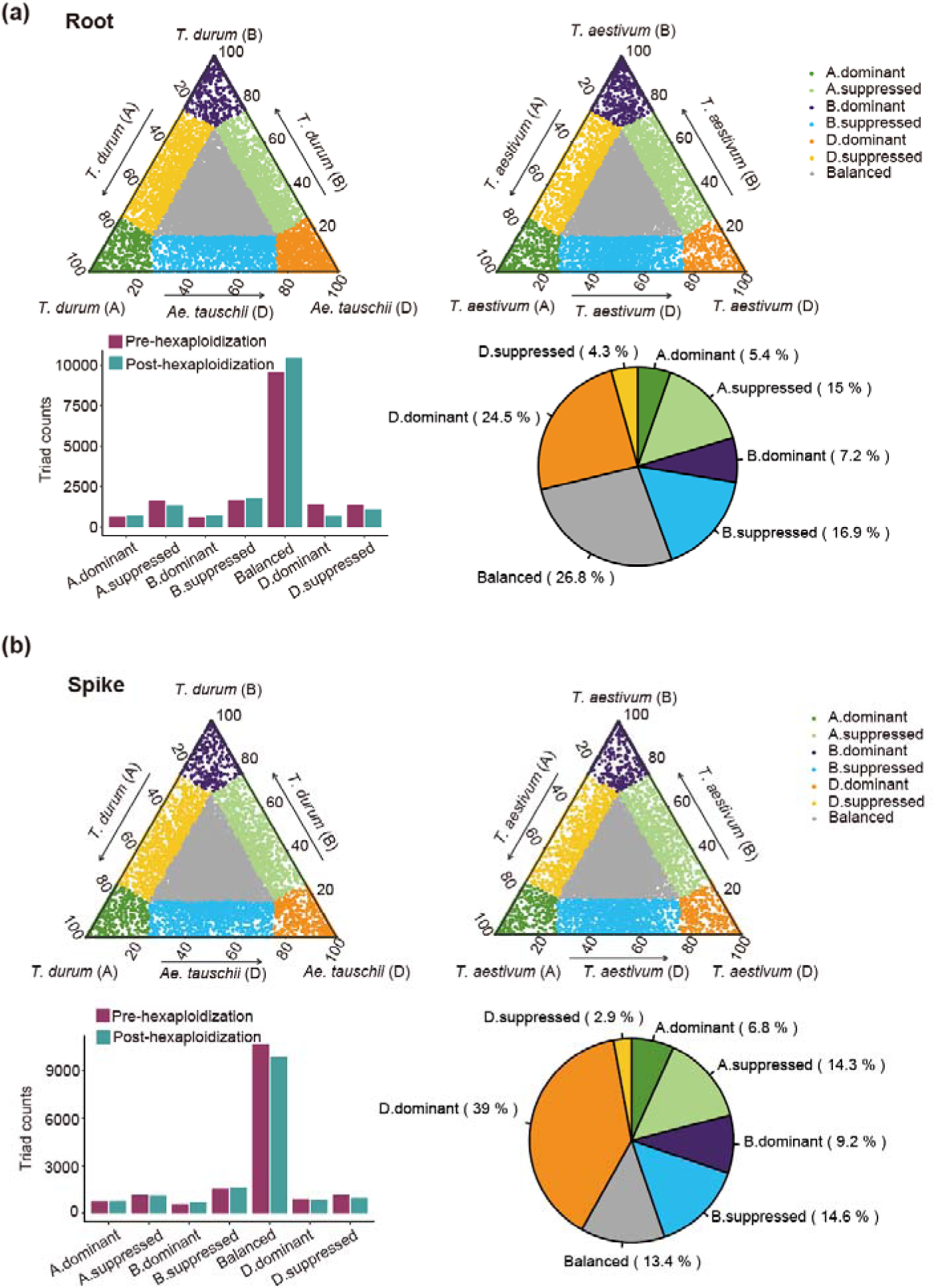
Expression dynamics of orthologous and homoeologous genes during polyploidization of wheat in root and spike. (a-b) Distribution of expression of orthologous genes between *T. durum* and *Ae. tauschii* (left) and homoeologous genes in A, B, and D subgenomes of *T. aestivum* (right) in root (a) and spike (b). The lower left panel showing the counts of triads in seven categories pre- and post-hexaploidization. And the lower right panel showing the classification and percentage of triads in post-hexaploidization that were classified into D.dominant category in pre-hexaploidization.

**Fig. S7.**
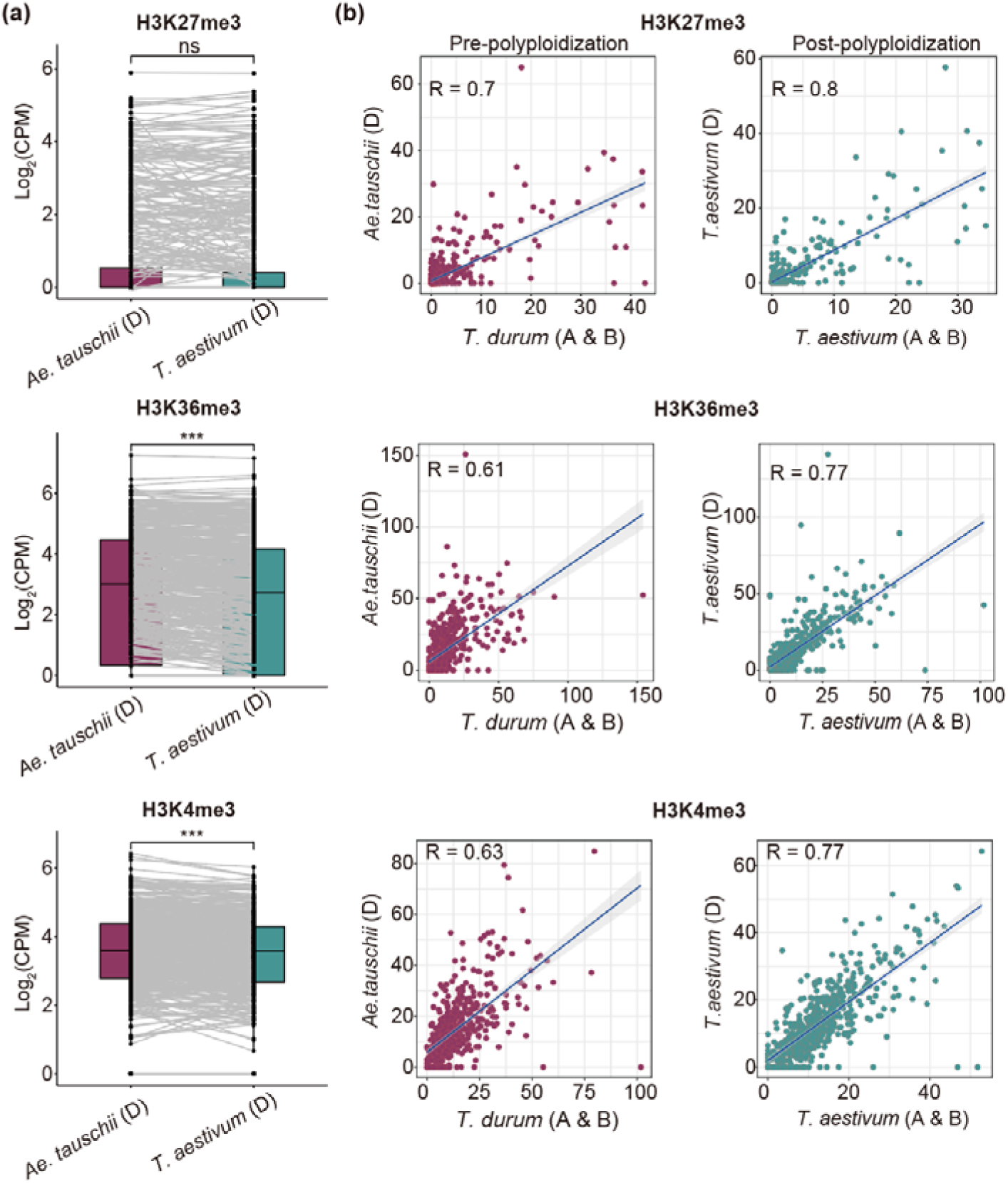
Change of histone modification in Dd-to-Db genes during polyploidization. (a) Comparison of H3K27me3, H3K36me3 and H3K4me3 signals of orthologs between *Ae. tauschii* and D subgenome in *T. aestivum*, which were classified into D.dominant to balanced categories before and after hexaploidization. Statistical significance was assessed by Student’s *t*-test. ***, *P* ≤ 0.001; ns, no significant difference. (b) Correlation of H3K27me3, H3K36me3 and H3K4me3 signals of orthologs between *Ae. tauschii* and *T. durum* (left) and homoeologs between D and A & B subgenomes in *T. aestivum* (right). The blue line denotes the line of best fit.

**Fig. S8.**
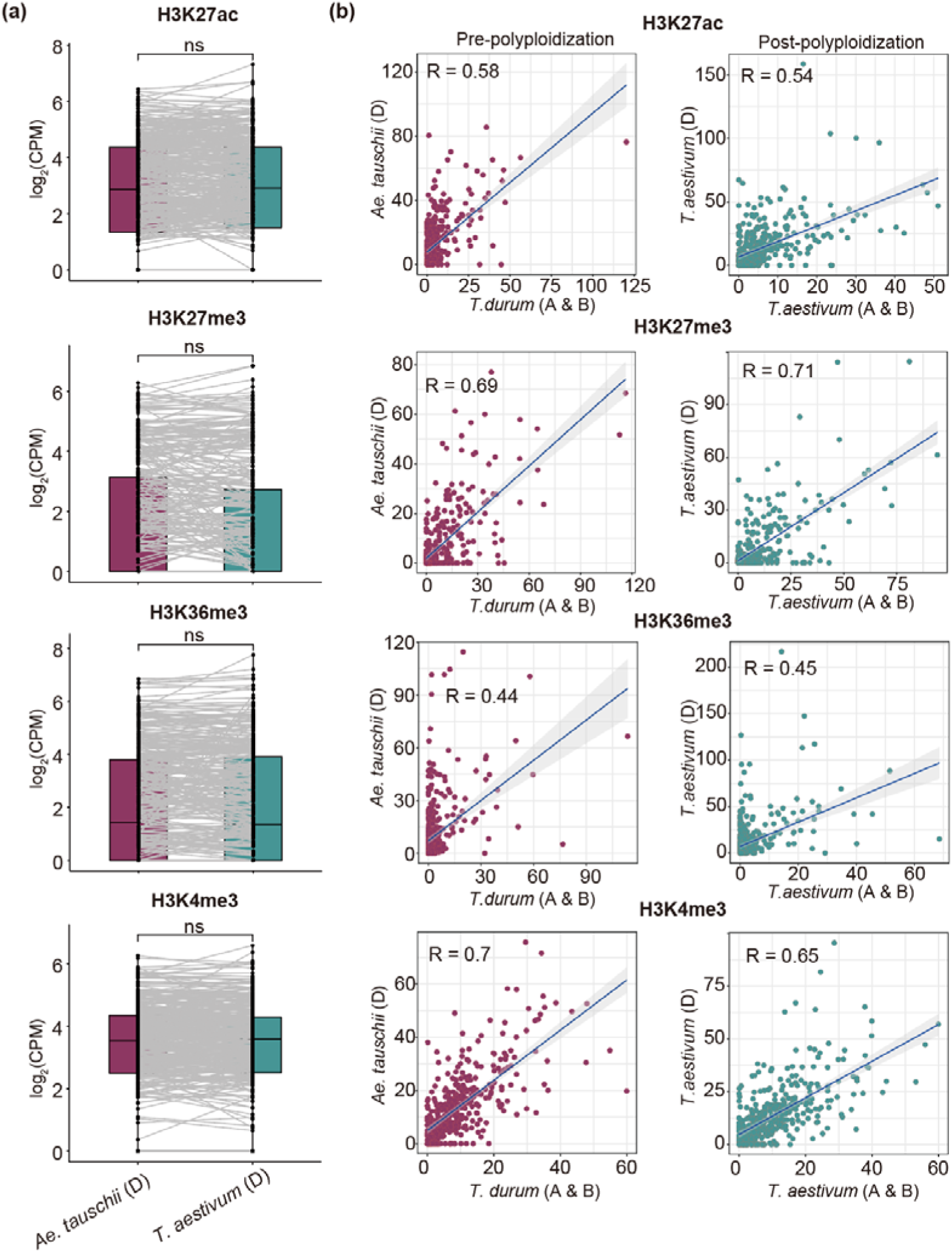
Change of histone modification in Dd-to-Dd genes during polyploidization. (a) Comparison of H3K27ac, H3K27me3, H3K36me3 and H3K4me3 signals of orthologs between *Ae. tauschii* and D subgenome in *T. aestivum*, which were classified into D.dominant to D.dominant categories before and after hexaploidization. Statistical significance was assessed by Student’s *t*-test. ***, *P* ≤ 0.001; ns, no significant difference. (b) Correlation of H3K27ac, H3K27me3, H3K36me3 and H3K4me3 signals of orthologs between *Ae. tauschii* and *T. durum* (left) and homoeologs between D and A & B subgenomes in *T*. *aestivum* (right), which were classified into D.dominant to D.dominant categories before and after hexaploidization. The blue line denotes the line of best fit.

**Fig. S9.**
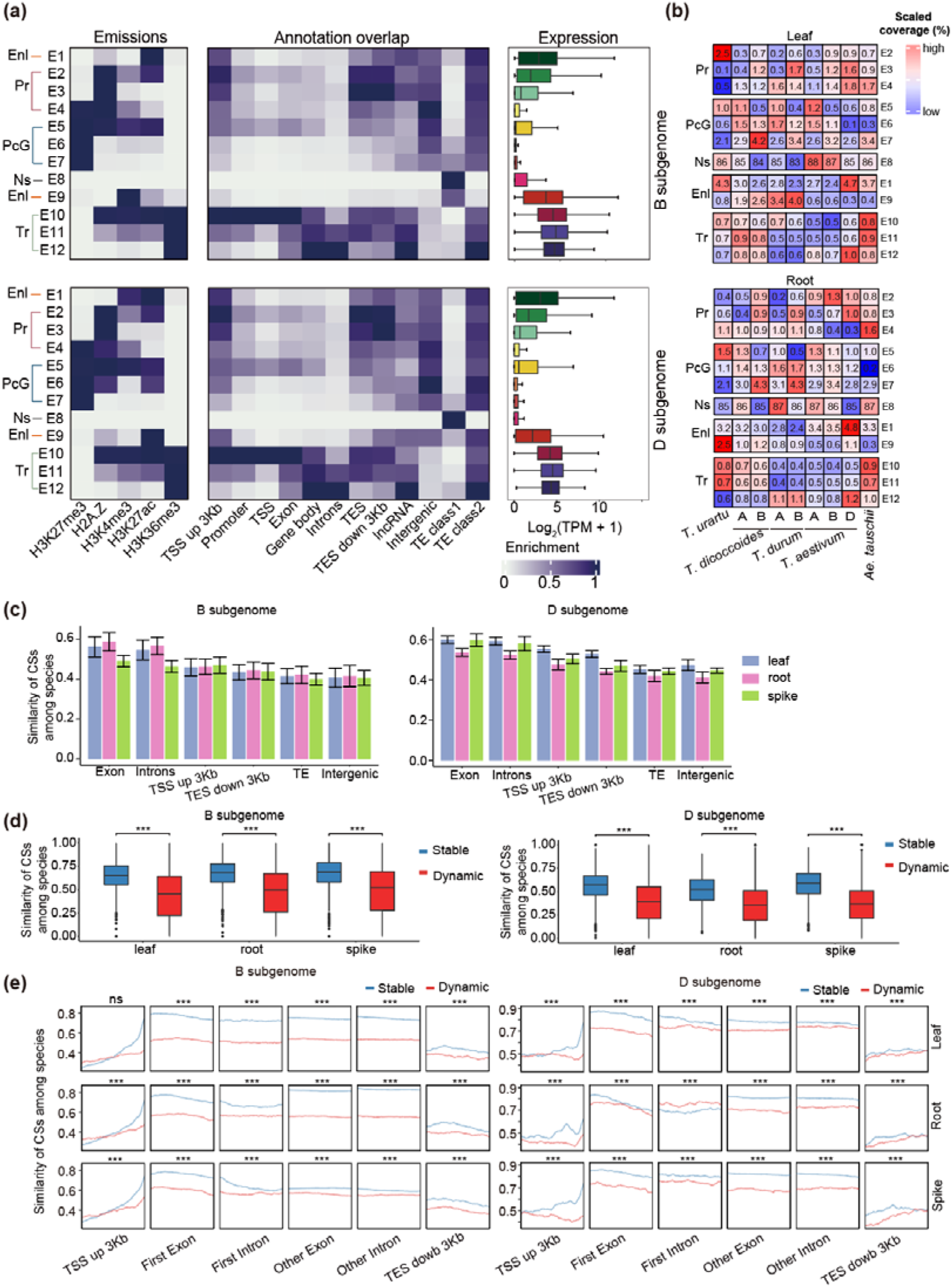
Chromatin states dynamic during the wheat speciation and the association with gene expression. (a) Definitions and overview of chromatin state annotations and epigenome data in B and D subgenome including emission probabilities for individual histone marks, and fold enrichments of chromatin states for the various types of genomic annotations and gene expression. (b) Comparison of the proportions of different CSs between the A, B and D subgenomes across five species in leaf and root. (c) Similarity of CSs among species in different genomic regions across leaf, root and spike in B and D subgenome. (d) Comparison of the similarity of CSs for stable and dynamic genes across leaf, root, and spike in B and D subgenome. Statistical significance was assessed by Student’s *t*-test. ***, *P* ≤ 0.001. (e) Distribution of similarity of CSs across gene body and flanking regions between stable and dynamic genes grouped by different transcription in B and D subgenome. Statistical significance was assessed by Student’s *t*-test. ***, *P* ≤ 0.001.

**Fig. S10.**
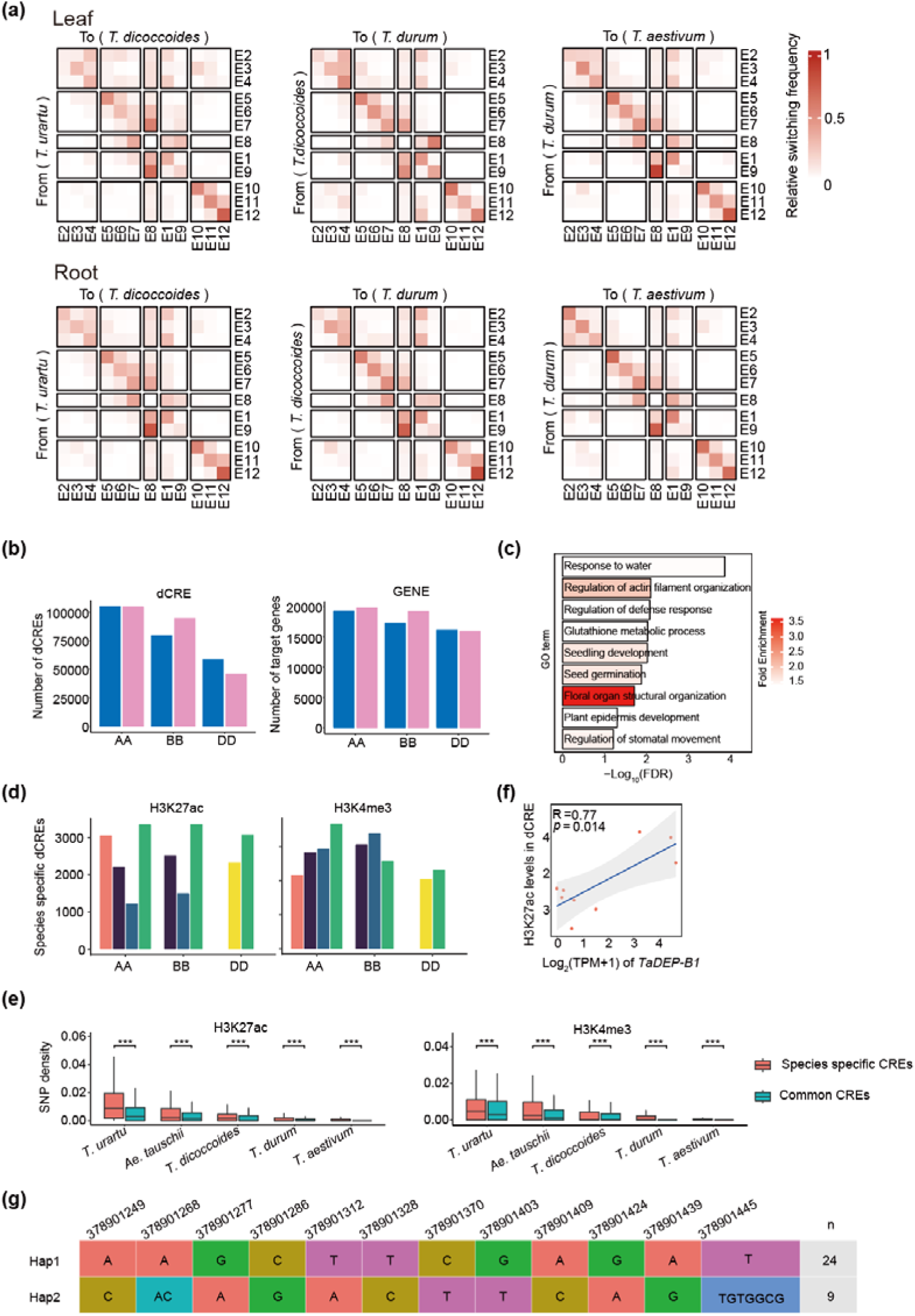
Dynamics of H3K27ac- and H3K4me3-mediated distal regulation during wheat speciation. (a) The switches of chromatin state during the polyploidization and evolution of wheat (from *T.urartu* to *T. dicoccoides*, *T. dicocccoides* to *T. durum*, and *T. durum to T. aestivum*). The color scale indicates the switching frequency. Ratio of the observed probability that a region switches from one CS (row) to another (column), The transition from Ns to Ns was removed. (b) The number of distal CREs mediated by H3K27ac and H3K4me3, and their target genes. (c) GO enrichment for H3K27ac and H3K4me3-mediated dCRE targeted genes. (d) The number of species-specific distal CREs mediated by H3K27ac and H3K4me3. (e) Species-specific SNP density in species-specific H3K27ac and H3K4me3-mediated dCREs across five species. Statistical significance was assessed by Student’s *t*-test. ***, *P* ≤ 0.001. (f) Correlation between expression levels of *TaDEP-B1* and H3K27ac levels of dCRE. The black line denotes the line of best fit. (g) Two typical haplotypes (Hap1 and Hap2) of *TaDEP-B1* were detected based on the resequencing data.

**Fig. S11.**
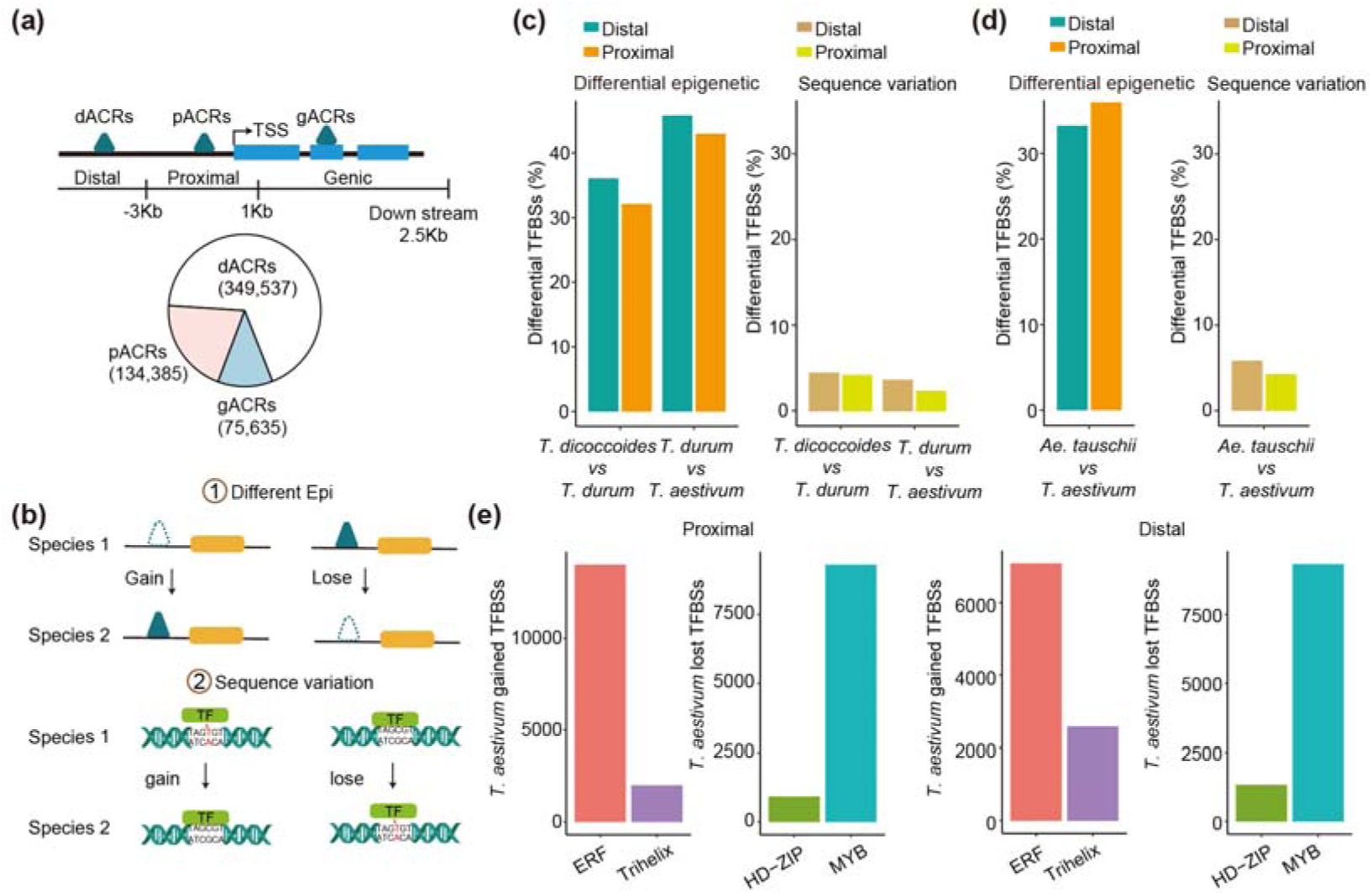
ATAC profile and changes of TFBSs during wheat speciation. (a) Subdividing ACRs based on distribution pattern around genes, and the number of dACRs, pACRs and gACRs. (b) Model of TFBSs gained or lost due to changed epigenetic modifications or sequence variations. (c) Ratio of changed TFBSs due to the alterations of epigenetic modifications or sequence variations across different stages in B subgenome. (d) Ratio of changed TFBSs due to the alterations of epigenetic modifications or sequence variations across different stages in D subgenome. (e) Proximal- and distal-specific loss of HD-ZIP and MYB binding sites and specific gain of ERF and Trihelix binding sites in *T.aestivum*.

**Fig. S12.**
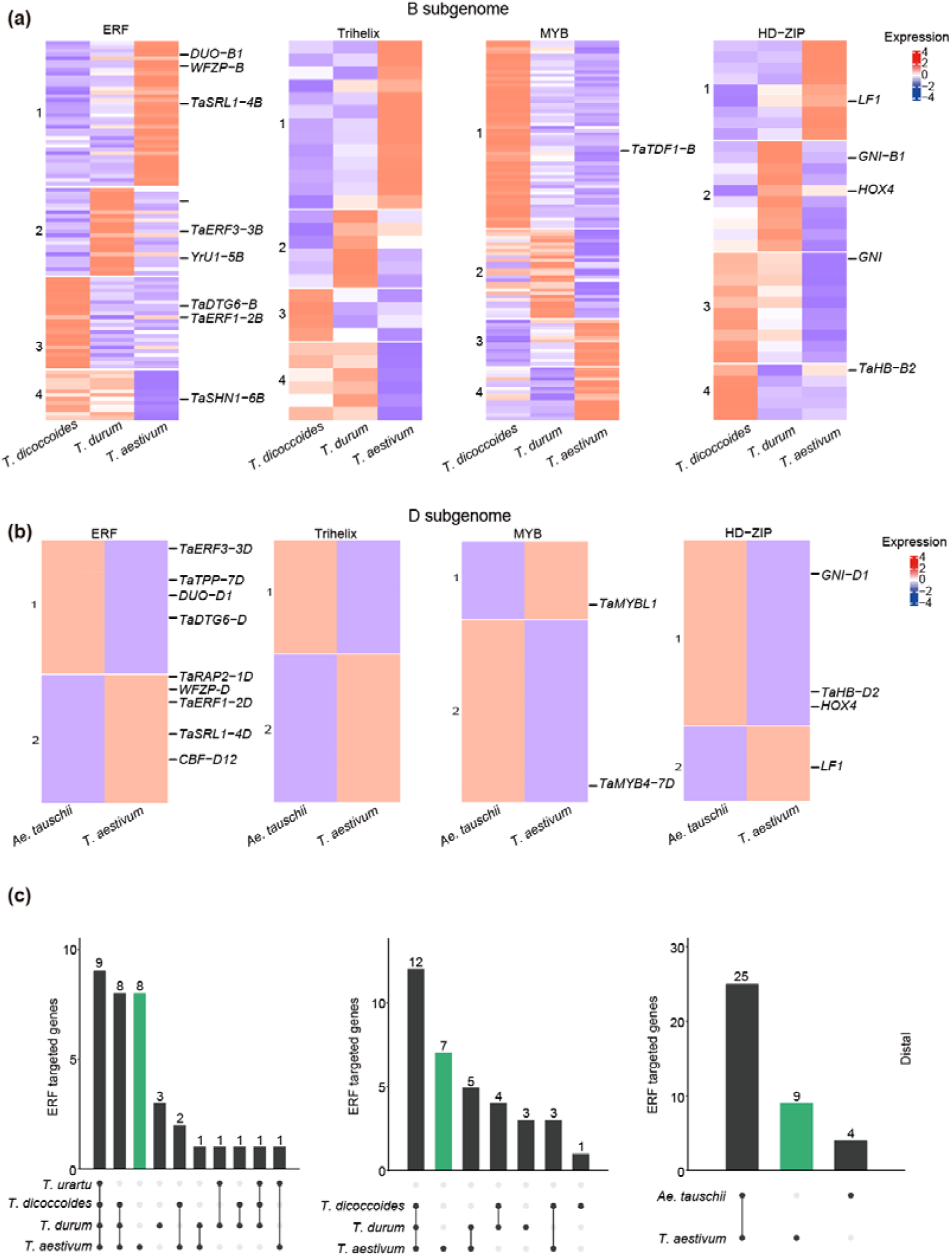
Expression levels of spike-related genes targeted TFs belong to ERF, Trihelix, MYB, HD-ZIP families during wheat speciation. (a-b) Relatively expression levels of ERF, Trihelix, MYB, HD-ZIP family TF sorted by *k*-means clustering across the four species of spike in B subgenome (a) or D subgenome (b). Known gene were labeled. (c) Overlap of spike-related genes (distal) targeted by ERF TFs between different species in A, B, and D subgenomes. The dots at the bottom represent the types of intersections among each species layer.

**Table S1** Sequencing and mapping statistics of CUT&Tag, ATAC-seq and RNA-seq data in *T. aestivum* and its tetraploid and diploid relatives.

**Table S2** Numbers of peaks detected for histone marks and chromatin accessibility in *T. aestivum* and its tetraploid and diploid relatives.

**Table S3** The coefficient of expression variations (CEV) for genes within each tissue across species in A, B and D subgenome.

**Table S4** Homoeolog expression bias across leaf, root, spike tissues in pre-polyploidization and post-polyploidization species.

**Table S5** Statistics and annotation of 12 chromatin states detected in *T. aestivum* and its tetraploid and diploid relatives.

**Table S6** Transition of 12 chromatin states among different species.

**Table S7** H3K27ac and H3K4me3 mediated dCRE-gene pairs.

**Table S8** Known spike-related genes containing ERF binding sites.

**Table S9** Antibodies and primers used in this study.

